# Body condition as an indicator of long-term ecosystem change: evidence from common dolphins

**DOI:** 10.64898/2026.09.08.750080

**Authors:** Marie A.C. Petitguyot, Juliette Champsaur, Alberto Hernandez-Gonzalez, Alfredo Lopez, Pablo Covelo, Jose Martínez-Cedeira, Xabier Pin, Mónica González, Uxía Vázquez, Graham J. Pierce

## Abstract

As climate change continues to disrupt marine ecosystems, reliable body condition indices (BCIs) for sentinel megafauna species such as dolphins will become increasingly important tools for long-term ecosystem monitoring. We developed novel BCIs from routinely collected morphometric measurements and applied them to a 22-year dataset (1998–2019) of 597 stranded common dolphins (*Delphinus delphis*) from Northwest Spain. BCIs were calculated as residuals from Generalized Additive Models relating body length to girth and blubber thickness measurements, thereby controlling for body size effects. The indices were first validated against visually assessed nutritional status and then used to investigate long-term trends and potential environmental and trophic drivers of body condition. The girth-based BCI outperformed the blubber thickness-based index in predicting nutritional status. The indices differed in their sensitivity to confounding variables. Both indices revealed an increase in male body condition until 2009–2010 followed by a sustained decline through 2019, whereas no long-term trend was detected in females. By integrating data on climate variability (North Atlantic Oscillation), oceanographic conditions (upwelling intensity, sea surface temperature, chlorophyll-a), prey availability, and prey quality, we found evidence that changes in body condition was associated with climate- and oceanography-driven changes in prey communities and reductions in the quality of prey available to dolphins. Our findings suggest that body condition in common dolphins integrates signals across multiple components of marine ecosystems and can serve as a sensitive indicator of ecosystem change. Given growing reports of declining body condition in marine predators worldwide, the approach presented here provides a practical and transferable framework for long-term ecosystem monitoring using existing stranding network datasets.

## 1. Introduction

Over recent decades, climate change and other anthropogenic stressors, such as fishing, shipping, coastal development, pollution, and non-indigenous species introductions, have reshaped marine ecosystems worldwide (Gissi et al., 2021; Ducklow et al., 2022), including the Northeast Atlantic (McQuatters-Gollop et al., 2022). Such ecosystem-level changes have their roots in lethal and sub-lethal impacts on individual animals. As well as having obvious implications for individual animal condition, health and welfare, such effects can provide an advance warning of changes at population, community and ecosystem levels. As such, indicators of individual condition and health of individuals of megafauna species have the potential to provide an important tool to monitor the impacts of climate change and other threats.

Body condition has been widely used as a proxy for nutritional status, and is by extension a measure of individual fitness, survival, and reproductive success (Castrillon and Bengtson Nash, 2020). The body condition of an animal integrates information on energy reserves (particularly lipid stores in blubber and muscle, which are known to vary in response to prey availability), reproductive status, and environmental stressors (e.g., Heide-Jørgensen et al., 2011; IJsseldijk et al., 2021). Various methods have been developed to assess the body condition of dead and live cetaceans, including the use of body condition indices (BCIs) derived from morphometric measurements (Castrillon and Bengtson Nash, 2020). Assessing the body condition of dead stranded individuals, rather than free-ranging animals, often has the advantage that there are multi-decadal datasets available, enabling the evaluation of long-term trends.

Since the late 1990s, the waters off Northwest Spain, particularly in Galicia, have experienced profound oceanographic and ecological changes. The region has shifted from cooler, drier conditions to warmer, wetter regimes, with altered wind patterns affecting upwelling (Bode et al., 2009, 2020), rising sea surface temperatures (Biguino et al., 2023; Piedracoba et al., 2024), and increasing ocean acidification (Padin et al., 2020). These changes have been accompanied by ecosystem regime shifts, including alterations in plankton and fish assemblages (Bode et al., 2009, 2020; Cabrero et al., 2019; Bañón et al., 2024), as well as declines in kelp forests, shellfish production, and changes in seabird dynamics (Voerman et al., 2013; Martínez-Abraín et al., 2019, 2023; Padin et al., 2024). Such ecosystem reorganization can cascade to upper trophic levels such as cetaceans, and potentially alter their foraging, physiology, and health (Ruiz-Cooley et al., 2017; Hildebrand et al., 2024), yet these effects remain largely unstudied in the Iberian Peninsula region.

The common dolphin (*Delphinus delphis*) is a widely distributed small delphinid inhabiting the Galician shelf and adjacent offshore waters (Gilles et al., 2023), and a generalist apex predator that feeds on a diverse range of small pelagic and mesopelagic fishes and squids, showing a preference for high-energy prey (Spitz et al., 2010; Santos et al., 2013; Marçalo et al., 2018; Faure et al., 2025; Hernandez-Gonzalez, et al., submitted). As top predators that occupy high trophic positions, common dolphins are ideal bioindicators to track climate and ecosystems changes (Ruiz-Cooley et al., 2017; Hazen et al., 2019). Evidence from stomach content analysis conducted on common dolphins stranded in Galicia indicates that they have experienced temporal shifts in their trophic ecology, likely driven by changes in prey availability (Santos et al., 2013; Hernandez-Gonzalez, et al., submitted). For example, the consumption of sardines decreased over time, while the consumption of hakes increased. Such changes in prey availability may influence not only diet composition but also prey quality and hence energetic intake, with potential consequences for individual condition and population dynamics (Gallagher et al., 2022). However, the effects of these ecological changes on the nutritional health of common dolphins in the region remain unclear, highlighting the need for further investigation.

In dead stranded cetaceans, measurements available to compute BCIs comprise the body length and various girth and blubber thickness measures, and sometimes the weight of the animal. Based on these measurements, the body mass index (BMI = body mass/body length^2^) is currently considered one of the most reliable indicators of body condition in stranded cetaceans (Kershaw et al., 2017; Castrillon and Bengtson Nash, 2020). However, weighing carcasses is not always feasible, as it depends on the logistical capacity and available resources of the stranding networks in charge of collecting dead animals. Consequently, body mass is recorded less frequently than girth measurements, blubber thickness, or nutritional condition codes (NCC) which are assigned on visual examination of the carcass (Petitguyot et al., 2025). As such, girth and blubber thickness measurements may be more accessible measures to compute BCIs.

Although stranding networks routinely collect such information, heterogeneity in data collection protocols among networks can complicate comparisons across regions and over time (Petitguyot et al., 2025). Not all networks systematically collect the same morphometric measurements. For example, current post-mortem examination protocols recommend measuring girth anterior to the dorsal fin (IJsseldijk et al., 2019), but stranding networks have historically collected a variety of girth measurements. These measurements have not yet been standardized, partly because their relevance as indicators of nutritional status remains poorly understood. Similarly, while standardized NCCs based on blubber and muscle assessment have been proposed as a five-category classification (IJsseldijk et al., 2019), their implementation varies widely, as some networks currently use a three- or two-category classification. Moreover, the assignment of NCCs is inherently subjective, as it depends not only on the examiner’s experience but also on the types of animals they routinely assess. In some regions, strandings may predominantly involve individuals in either very good or very poor condition, limiting opportunities for comparison and making it difficult to evaluate an animal’s status against a representative ‘normal’ nutritional baseline.

The choice of an appropriate BCI also depends on the assumptions underlying its calculation and the specific objectives for which it is intended. For example, some indices may be selected to best represent the NCC given during a visual examination of the carcass (e.g., Albrecht et al., 2024) or the body mass (e.g., Stepien et al., 2023), and others to be independent of body length and to discriminate among causes of death, age classes and reproduction seasons (e.g., Kershaw et al., 2017). Various BCIs have been proposed for use in dolphins and porpoises. For example, in captive harbour porpoises (*Phocoena phocoena*) maintained in semi-enclosed facilities, Stepien et al. (2023) found that the girth measurements taken immediately in front of the dorsal fin, and behind and in front of the pectoral fins were the most reliable indicators of body condition. In contrast, Albrecht et al. (2024) reported that ventral blubber thickness was the best predictor of nutritional status in dead stranded common dolphins from the Celtic Seas ecoregion. The average blubber thickness is used as an indicator of nutritional status in seals in the HELCOM area (i.e., area covered by the Baltic Marine Environment Protection Commission (Helsinki Commission); HELCOM, 2018). The use of blubber thickness as an indicator of nutritional status has been questioned, given that, in small cetaceans, it not only serves as an energy reserve but also plays important physiological roles, such as thermal insulation, and thus may not fully reflect nutritional status alone (Derous et al., 2020). Further research is needed to evaluate whether girth, blubber thickness and weight measurements actually provide meaningful indicators of nutritional status and overall health. In addition, girths and blubber thicknesses are often related to body length. One approach to account for this relationship is to use residuals from regressions between girth or blubber thickness and body length. This method has been widely applied to derive BCIs from girth – length, blubber thickness – length, and body mass – length (e.g., Read, 1990; Haug et al., 2002; Gómez-Campos et al., 2011; Kershaw et al., 2017) relationships using Ordinary Least Squares (OLS) regression. However, OLS assumes linear relationships between variables, which may not always be the case (Green, 2001). Relationships may also be specific to species, population, sex and age-class or simply differ between different data sets (Labocha et al., 2014; Falk et al., 2017). Generalized Additive Models (GAMs) can provide a more flexible alternative to OLS, by accommodating non-linear relationships, as used for body mass – length relationships in fish (e.g., Pierce et al., 2018). Furthermore, by quantifying body condition relative to the population average, residuals from the population-level model can be used to define thresholds for classifying and interpreting population-level body condition.

In this study, we used body condition as a proxy for nutritional status to examine whether the major climate regime shifts and associated ecosystem changes over recent decades in Galicia have influenced the nutritional health of common dolphins. We explored spatio-temporal variation in the body condition of common dolphins stranded along the Galician coast during 1998-2019, in relation to various potential drivers. As the weight of animals is not systematically recorded in Galicia, we computed five body length-independent BCIs derived from available morphometric measurements collected by the Galician stranding network, including two girth measurements and three blubber thickness measurements. BCIs were computed as residuals from GAMs relating body length to each of the five measurements, thereby accounting for body-length effects. We hypothesized that girth-based and blubber thickness-based BCIs would behave differently and that both should therefore be evaluated. Accordingly, our first objective was to identify the most informative girth-based and blubber thickness-based BCIs for predicting the NCC visually attributed during necropsy.

We then used these two BCIs to (1) assess spatio-temporal trends in the body condition of male and female common dolphins, and evaluate the influence of potential confounding factors (season, decomposition state) using GAMs, ensuring that the indices could reliably differentiate the animal’s health status (per cause of death assessed during necropsy); and (2) investigate potential trophic and environmental drivers of body condition, including proxies for prey quality (average energy density of prey per stomach (kJ g^-1^), based on stomach content analysis), prey availability (Spawning Stock Biomass of some of the main prey species on common dolphins in Galicia), primary production (surface chlorophyll-a concentrations), and environmental variability (sea surface temperature, local upwelling intensity, North Atlantic Oscillation) using temporal trends analyses and piecewise Structural Equation Modelling.

By integrating morphometric, ecological, and environmental data, this work aims to (i) develop reliable BCIs adapted to common dolphins stranded in Galicia (and potentially more broadly applicable), (ii) better understand what factors influence body condition, and (iii) provide insight into how ecosystem changes in Galician waters may have influenced the nutritional status of this key marine predator.

## 2. Material and methods

### 2.1. Study area and data collection on stranded dolphins

The Galician stranding network, coordinated by the non-governmental organization Coordinadora para o Estudio dos Mamíferos Mariños (CEMMA), has systematically responded to cetacean strandings in Galicia since 1990. The network conducts necropsies on stranded individuals and on bycaught animals obtained through direct collaboration with fishers, following the standardized protocol of the European Cetacean Society (Kuiken and García-Hartmann, 1991). It records a comprehensive set of data and collects various biological samples during post-mortem examinations.

Here, we used a dataset of 597 common dolphins mainly stranded animals along Galician shores but also some bycaught and handed in by fishers between 1998-2019, thus spanning a 22-year time period (Figure 1.a). The dataset includes morphometric measurements of the total body length measured as the distance from the tip of the rostrum to the notch in the tail fluke (in cm), the girth around the body taken behind the pectoral fins (cm), the girth taken at the post-nuchal depression (cm), and the dorsal, lateral, and ventral blubber thickness (mm) (Figure 1.c). Not all measurements were available for every dolphin and, of the total sample, 341 individuals had a complete set of morphometric measurements. The information collected also includes the date and location of stranding, sex, the NCC (classified as ‘good’, ‘moderate’, or ‘poor/very poor’ based on a visual assessment of the blubber and muscle condition, following standard criteria adapted from Kuiken and García-Hartmann, 1991; see Table S1 for definitions), stage of carcass decomposition (DCC; on a scale from 1 to 5: DCC1 = stranded alive and died immediately after; DCC2 = recently dead and extremely fresh; DCC3 = moderate decomposition; DCC4 = advanced decomposition; DCC5 = mummified or skeletal remains), and cause of death (COD) with respect to bycatch (i.e., when COD could be diagnosed, whether or not there was evidence of bycatch).

**Figure 1.**
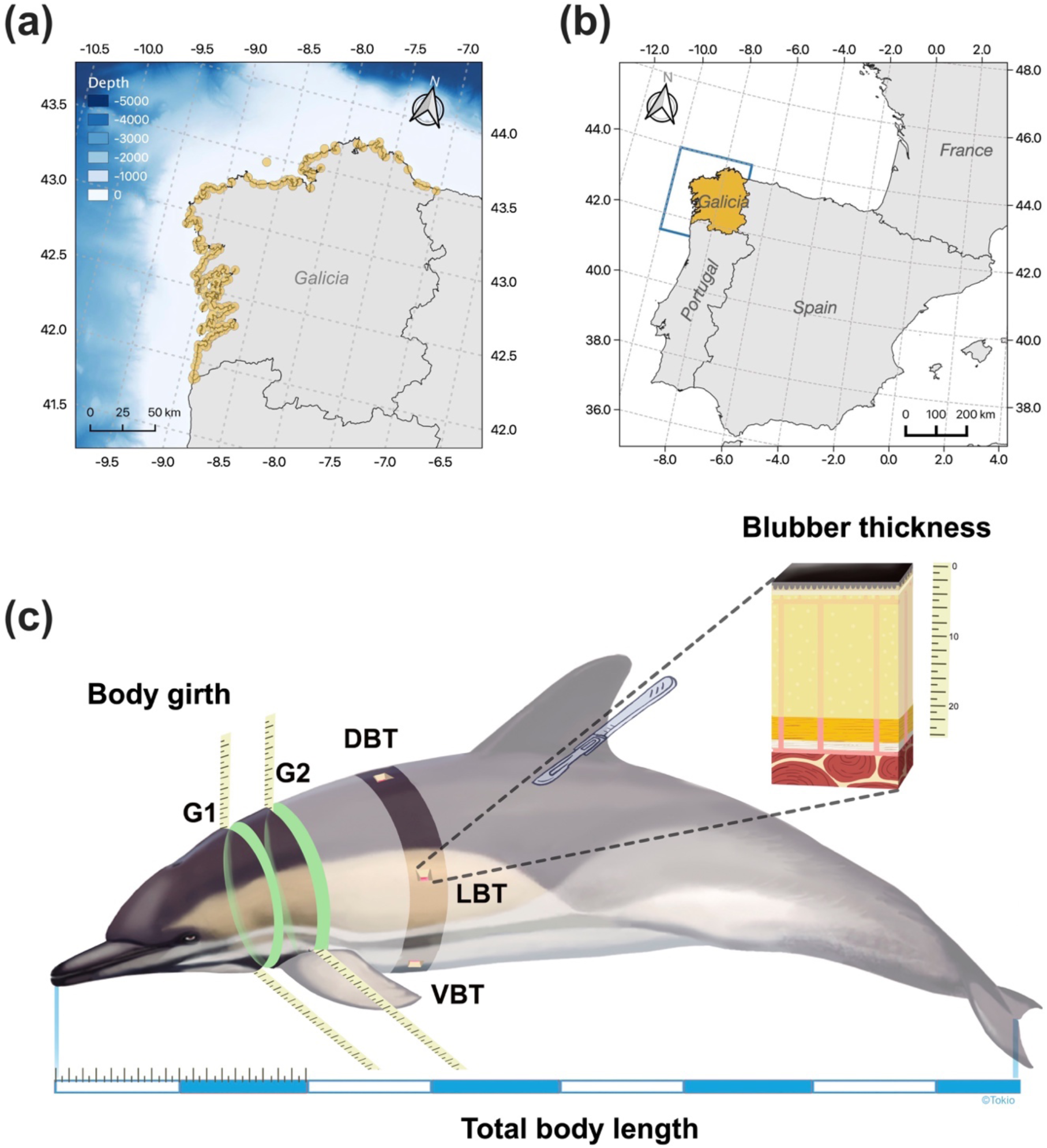
(a) Map of the study area showing the location of the common dolphins stranded along the coast or bycaught at sea used in this study. (b) Environmental data obtained from the Copernicus Marine Environment Monitoring Service cover the study domain (41.5°/44° N, −10°/-7° W), highlighted by the blue square. Maps were produced using QGIS 3.8.1. (c) Body measurements collected during carcass examination by the Galician stranding network. G1: girth taken at the post-nuchal depression; G2: girth taken behind the pectoral fins; DBT: dorsal blubber thickness; LBT: lateral blubber thickness; VBT: ventral blubber thickness; total body length. Illustration by Alfredo Lopez - Tokio illustration ©.

### 2.2. Proxy for prey quality

The stomach contents of 448 common dolphins stranded in Galicia between 1998-2018 were analyzed by Santos et al. (2013) and Hernandez-Gonzalez et al. (submitted). Of these, 276 individuals had at least one measurement recorded in our morphometrics dataset. The identification of prey in the diet was carried out following methods described in Santos et al. (2013) and Hernandez-Gonzalez et al. (2024). Identification of fish, cephalopods, crustaceans and molluscs was done to the lowest possible taxonomic level using both published guides (e.g., Clarke, 1986; Härkönen, 1986; Watt et al., 1997) and reference material. The estimated biomass of prey species contained in each stomach was converted into energy density values (kJ g^-1^) using the fish energy density data from Spitz et al. (2010).

### 2.3. Proxy for prey availability

We used the Spawning Stock Biomass (SSB; in tonnes) of the main commercially important prey species of common dolphins in Galicia as proxies for prey availability, as per Santos et al. (2013) and Hernandez-Gonzalez et al. (submitted) in ICES areas 27.8c and 27.9a covering Galician waters, i.e., blue whiting *Micromesistius poutassou*, Iberian sardine *Sardina pilchardus*, European hake *Merluccius merluccius*, Atlantic horse mackerel *Trachurus trachurus*. For some of these species, SSB estimates were not available specifically for areas 27.8c and 27.9a and we used the values reported for larger areas. The SSBs of (1) blue whiting (WHB) in ICES Subarea 27.1–9, 12 and 14, (2) Iberian sardine (PIL) in ICES Subarea 27. 8.c and 9.a, (3) European hake (HKE) in ICES Subarea 27. 8.c and 9.a, and Atlantic horse mackerel in (4) ICES Subarea 27.8 and divisions 2.a, 3.a, 4.a, 5.b, 6.a, 7.a–c, e–k (HOMne) and (5) ICES Subarea 27.9.a (HOM9.a), were extracted from the ICES Stock Assessment Database, and the 2024 annual reports of the Working Group on southern horse mackerel, anchovy and sardine (WGHANSA) and the Working Group on widely distributed stocks (WGWIDE) (ICES, 2024b,c). To provide a comprehensive estimate of total Atlantic horse mackerel availability, the SSBs of HOMne and HOM9.a were integrated into a single representative variable (HOM). Annual SSB for HOM was calculated by summing the mean values of the HOMne and HOM9a stocks. Combined uncertainty was determined by propagating standard errors 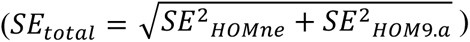, with standard errors derived from the reported 95% confidence intervals.

### 2.4. Environmental and climatic data

Sea surface temperature (SST, °C), surface chlorophyll-a concentrations (CHL, mg m^-3^), upwelling intensity (UI), and the North Atlantic Oscillation (NAO) were selected as local and large-scale environmental and climatic variables likely to provide insight into physiological and trophodynamic processes in top predators (e.g., Young et al., 2015; Ruiz-Cooley et al., 2017; Lambert et al., 2022). CHL, commonly used as proxy for prey availability (e.g., Lambert et al., 2022), and SST were obtained from the Copernicus Marine Environment Monitoring Service (https://marine.copernicus.eu/access-data). Specifically, the CHL was sourced from the Copernicus Iberia-Biscay-Ireland biogeochemical reanalysis (Atlantic-Iberian Biscay Irish-Ocean BioGeoChemistry, https://doi.org/10.48670/moi-00028), while SST was retrieved from the corresponding physical reanalysis product (https://doi.org/10.48670/moi-00029). Daily data spanning 1998–2019 for the study area (latitude 41.5° - 44°, longitude −10° - −7°; Figure 1.b) were extracted at a 0.083° × 0.083° spatial resolution and averaged by month to obtain monthly means.

The UI (m^3^ s^-1^ km^-1^) are based on calculation of the Ekman transport from surface winds by the Instituto Español de Oceanografía (http://www.indicedeafloramiento.ieo.es/). Each value refers in a cell of 1°×1° centered at the position 43°N 11°W (considered as representative of the upwelling intensity on the west coast of Galicia), using data from atmospheric pressure at sea level derived from the WXMAP model (González-Nuevo et al., 2014). Monthly UI means were extracted for the period 1997-2019. Positive values of this index indicate net upwelling periods when surface water is transported offshore, while negative values indicate an accumulation of surface water against the coast (downwelling). Daily values of the NAO were averaged to monthly means for the period 1997-2019 and were obtained from the National Oceanic and Atmospheric Administration (NOAA) (https://www.cpc.ncep.noaa.gov/).

### 2.5. Statistical analyses

#### 2.5.1. Computation of body condition indices

Data exploration following Zuur et al. (2010) indicated non-linear relationships between body condition measurements and total body length (Figure S1). Girth measurements in particular showed a cone-shaped dispersion pattern relative to body length, suggesting increasing variance with size (heteroscedasticity). Therefore, girth measurements were log-transformed before further analysis. We tested for sex-specific differences in the relationships between total body length and each morphometric measurement by comparing the Akaike’s Information Criterion (AIC) of two GAMs: one with a shared smooth of total body length and one allowing sex-specific smooths. As the AIC values differed by less than two units, we found no substantial support for sex-specific relationships and therefore did not compute BCIs separately for each sex (Table S2). Our objective was to derive a BCI that was independent of body length, allowing us to investigate spatio-temporal patterns without confounding effects from size. Given the strong relationship between body length and the morphometric measures in our dataset, we applied a residual-based approach. Specifically, we calculated five BCIs as residuals from GAMs that modeled girth and blubber thickness as functions of body length:

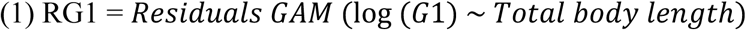

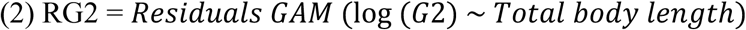

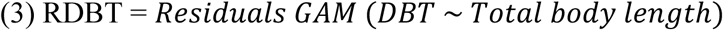

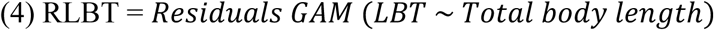

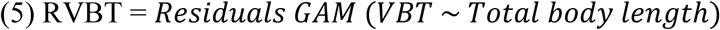

These residual-based indices describe the relative position of each individual in our dataset with respect to the expected morphometric-length relationship. Positive values indicate that an individual’s girth or blubber thickness exceeds the value predicted for its body length, whereas negative values indicate measurements below the predicted value. These deviations were interpreted as reflecting relatively higher and lower body condition, respectively.

#### 2.5.2. Selections of BCIs to predict nutritional condition

Our selection criteria were based on two main assumptions: (1) the BCI should accurately predict the NCC assigned during carcass examination, and (2) it should effectively discriminate between animals with differing health status.

In the absence of long-term pathological data to classify individuals by health status, we relied on the cause of death (COD) category recorded during post-mortem examinations, which consisted of whether or not the animals showed evidence of bycatch. Animals that died as a result of bycatch are assumed to be more representative of the free-ranging population and to have generally been in better body condition prior to death, even though they might have presented some health issues. In contrast, animals that did not die from bycatch are more likely to have experienced underlying health issues and are therefore assumed to have been in poorer body condition. Consequently, a robust BCI should be able to distinguish between these two groups.

We expected a priori that because the NCC is assigned through visual assessment of blubber and muscle condition, and likely reflects the overall external body shape more strongly than specific muscle or blubber characteristics, the girth-based BCIs would better represent the assigned NCC than the blubber thickness-based indices. We also assumed that girth-based BCIs and blubber thickness-based BCI would behave differently and should therefore be evaluated in a complementary manner. Thus, consistent with both assumptions, we aimed to select the best girth-based BCI and the best blubber thickness-based BCI for representing the NCC. To determine which of the five computed BCIs best predicted the NCC, we followed a three-step approach based on analyses restricted to a subset of animals with complete data for all five indices (see Figure 2 for details regarding datasets and sample size used in each research question).

**Figure 2.**
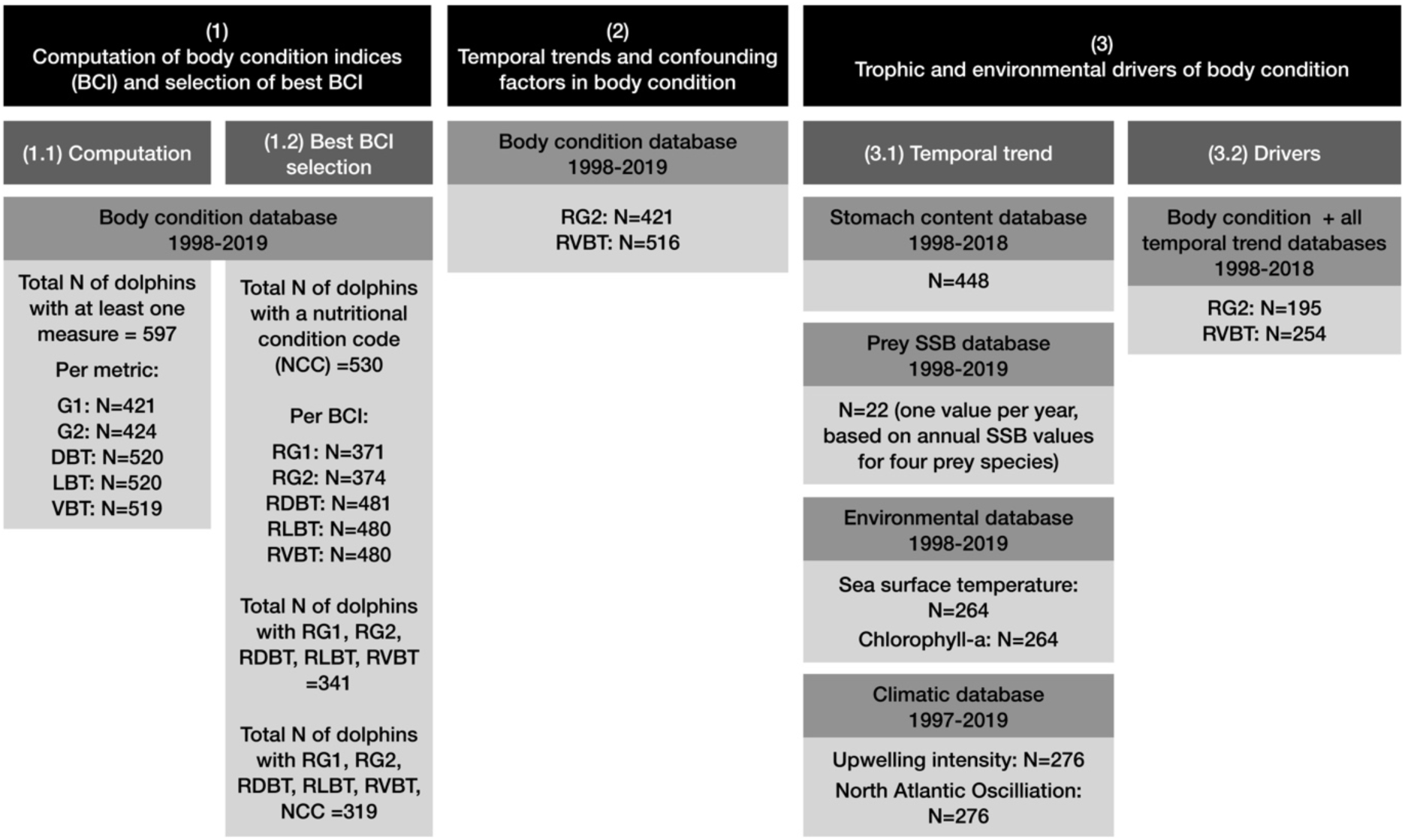
Overview of the research design, detailing databases, time periods, and sample sizes across specific research objectives.

We first assessed pairwise correlations among the five BCIs using Pearson’s correlation coefficients on a subset of animals with all 5 BCIs available (N=341). Correlation coefficients were calculated for each BCI pair, with statistical significance assessed at *α* = 0.05. Second, we conducted a principal component analysis (PCA) on the five BCIs to evaluate their multivariate structure and identify patterns of covariation among indices. The analysis was done on a subset of animals that had all 5 BCIs and the NCC available (N=319). All variables were mean-centered and scaled to unit variance prior to the analysis, to ensure that indices measured on different scales contributed equally. Due to relatively few individuals classified as having a ‘poor/very poor’ nutritional condition, animals categorized as ‘moderate’ and ‘poor/very poor’ were combined into a single ‘moderate/poor’ group for subsequent analyses. NCC was not included in the PCA computation, as the objective was to explore the intrinsic correlation structure of the BCIs (unsupervised analysis). However, the NCC (‘good’ vs. ‘moderate/poor’) was overlaid onto the PCA ordination to visually assess group separation along the principal component axes. To facilitate interpretation, the loading vectors of each BCI were plotted as arrows originating from the origin in the biplot. The direction of each arrow indicates the gradient along which that index increases in the multivariate space, while the length of the arrow reflects the strength of its contribution (i.e., magnitude of loading) to the principal components displayed. Indices with longer arrows and orientations aligned with separation between nutritional groups were interpreted as contributing most strongly to the observed NCC gradient.

Finally, we fitted five separate logistic generalized linear models (GLMs), each testing the relationship between a single BCI and the NCC, on separate subsets (e.g., RG1: N=371; see Figure 2). For each model, we estimated the P50 value to determine the threshold distinguishing ‘good’ versus ‘moderate/poor’ condition. Model performance was compared using AIC, and the best-performing models were selected as those that most accurately predicted the NCC.

#### 2.5.3. Temporal trends in body condition and confounding variables

The best ranked girth- and blubber thickness-based BCIs were subsequently used to evaluate spatio-temporal variation in body condition and the effects of potential confounding factors likely to influence the body condition of common dolphins. All data were explored extensively following Zuur et al. (2010) to inform model selection. As the data exploration revealed non-linear trends, GAMs with a Gaussian distribution were selected to assess variation in body condition. As measurements and information on confounding variables were not available for all animals, GAMs with the best girth-based BCI (RG2) were applied to 421 individuals, and those with the best blubber thickness-based BCI (RVBT) to 516 individuals (Figure 2).

Temporal variations were modelled using the year of stranding (continuous; numeric representation where each date is expressed as a fraction of the year), and the season (continuous; represented as the day of sampling within each year, modelled using a cyclic penalized cubic regression spline to account for its cyclical nature). Spatial structure was incorporated by introducing a north-south gradient by using the latitude of the stranding location (continuous).

We assessed differences between sexes (categorical; males; females). In addition, known females carrying a foetus were excluded from analyses using girth-based BCI to avoid biasing these values. The presence of a foetus may inflate the girth, particularly the one taken behind the pectoral fins, which is situated closest to the womb. However, it is possible that some pregnancies were not detected and as such some pregnant females could have been included in our analyses.

We assessed the effect of the decomposition stage of the carcass on body condition values (categorical; DCC1_2 = alive or recently dead and extremely fresh; DCC3_4 = moderate or advanced decomposition - individuals classified as DCC1 and DCC2, and as DDC3 and DCC4, were combined into a single category due to the small sample sizes in each group). As carcasses decompose, they progressively bloat and may leak blubber. These processes can distort soft tissues and consequently bias girth and blubber thickness measurements.

To ensure that the BCIs could reliably differentiate health status, the COD category which noted the presence or absence of bycatch indicators was added in the models (categorical; evidence of bycatch = Yes; no evidence of bycatch = No).

All GAMs were fitted using the *mgcv* package (Wood, 2011). We compared models using the Akaike Information Criterion (AIC; Akaike, 1987) to evaluate various combinations of interactions with sex in each of the smoother terms, and retained only the interaction between sex and year. Model selection was based on AIC, using a manual backward stepwise procedure. Model performance and assumptions were assessed through standard diagnostic and validation checks using the *DHARMa* package. Multi-collinearity was tested between non-linear predictors using the concurvity() function and basis functions (k) were checked using the gam.check() function.

Model effects were visualized with the *ggplot2* package (Wickham, 2016) based on fully marginalized predictions obtained with the *marginaleffects* package (Arel-Bundock et al., 2024). Rather than holding non-focal covariates at fixed mean or reference values, predictions were generated across grids spanning the observed values of all variables retained in the final GAMs. Predicted responses were then averaged over the remaining covariates to estimate the marginal effect of each focal predictor. This approach yields population-averaged model predictions on the original response scale while preserving the interpretation of BCI=0 as the reference condition.

#### 2.5.4. Trophic and environmental drivers of body condition

Data exploration revealed collinearity and auto-correlation within the four prey SSBs. To integrate the SSB data of WHB, PIL, HKE, and HOM into a single index of prey availability, we performed a Dynamic Factor Analysis (DFA) using the *MARSS* package (Holmes et al., 2025). This method models the auto-correlation inherent in biological time-series, where current biomass is dependent on previous states. DFA allowed for the extraction of a single latent trend representing the common trajectory of prey availability.

We first examined long-term trends in all trophic and environmental variables. We fitted GAMs with a Gaussian distribution to evaluate the relationships with year for the proxies of both prey quality (energy density per stomach stratified by sex; N=448) and prey availability (DFA index; N=22). To isolate long-term ecological trajectories from high-frequency noise and seasonal fluctuations, environmental time series of CHL (N=264), SST, (N=264), UI (N=276), and NAO (N=276) were subjected to Seasonal-Trend Decomposition using Loess (STL; Cleveland et al., 1990). This additive approach partitioned the raw monthly observations into seasonal, trend, and remainder components, allowing for a precise evaluation of the underlying de-seasonalized signals. Following decomposition, the trend components were analyzed for monotonic directions using the non-parametric Mann-Kendall (MK) test distribution (Mann, 1945; Kendall, 1975) with the *Kendall* package (McLeod, 2022) and Sen’s Slope estimator using the *trend* package (Pohlert, 2020), providing the rate of change per year. To investigate potential regime shifts during the study period, structural breakpoint analysis was applied to the STL trends using the Bai & Perron (2003) method and the *strucchange* package (Zeileis et al., 2002). The optimal number and timing of structural breaks were determined by minimizing the Bayesian Information Criterion. This two-step methodology ensured that detected shifts represented fundamental changes in the mean state of the ecosystem rather than transient seasonal anomalies, enabling a synchronized comparison of atmospheric forcing and biological response across the 1997/1998-2019 period.

To investigate the bottom-up drivers of dolphin body condition, we used a reduced subset of the data (RG2: N=195; RVBT: N=254; see Figure 2), due to information on energy density per stomach not being available for every dolphin. We conducted piecewise Structural Equation Modeling (pSEM) using the *piecewiseSEM* package (Lefcheck, 2016). This approach allowed for the integration of multiple hierarchical relationships into a single causal framework while accounting for the non-linearities common in ecological time series through Gaussian GAMs using the *mgcv* package. For each BCI subset, we constructed an *a-priori* pSEM hierarchical model representing four ecological paths as a bottom-up cascading effect: (sub-model 1) Atmospheric forcing, as the influence of the NAO decomposed trend (NAO_deco) on UI_deco; (sub-model 2) Primary productivity, as the combined effect of UI_deco and SST_deco on CHL_deco; (sub-model 3) Trophic intermediate level, as the response of the prey quantity (DFA index) to CHL_deco and UI_deco and SST_deco; (sub-model 4) Higher trophic level, as the effect of prey availability (DFA index) and prey quality (energy density) on dolphin body condition (RG2 or RVBT). Sex-specific smoothers were tested via AIC for all dietary predictors in sub-model 4. The interaction was retained only for the DFA index of the RG2 pSEM, as it significantly improved model fit and captured divergent responses between sexes; energy density variables were subsequently modeled without sex interaction. To account for biological delays in ecosystem response to large-scale climatic variables (UI and NAO), we compared competing hypotheses using AIC in sub-model 2 and 3: a Year^-0^ model (immediate ecological response) and Lagged models incorporating combinations of a year^-1^ delay in UI and NAO. The Year^-0^ model demonstrated superior fit and was retained for all subsequent analyses. We further investigated whether environmental and seasonal drivers exerted direct physiological or foraging stress on dolphins, rather than being strictly mediated through the food web. To do so, we used AIC to compare the *a-priori* hierarchical pSEM model against alternative structures containing direct paths from SST_deco, UI_deco, NAO_deco, and the day of the year (modelled using a cyclic penalized cubic regression spline to account for its cyclical nature), to dolphin body condition (RG2 and RBVT). Each sub-component of the pSEM was individually validated using the *DHARMa* package, and both multi-collinearity between non-linear predictors and basis functions (k) were checked.

## 3. Results

### 3.1. Selection of BCIs to predict nutritional condition

Pearson correlation coefficients showed that the two girth-based indices were strongly correlated with each other, as were the three blubber-based indices. In contrast, correlations between the girth- and blubber-based indices were only moderate (Figure 3.a), suggesting that these BCI do not respond in the same way to external factors.

**Figure 3.**
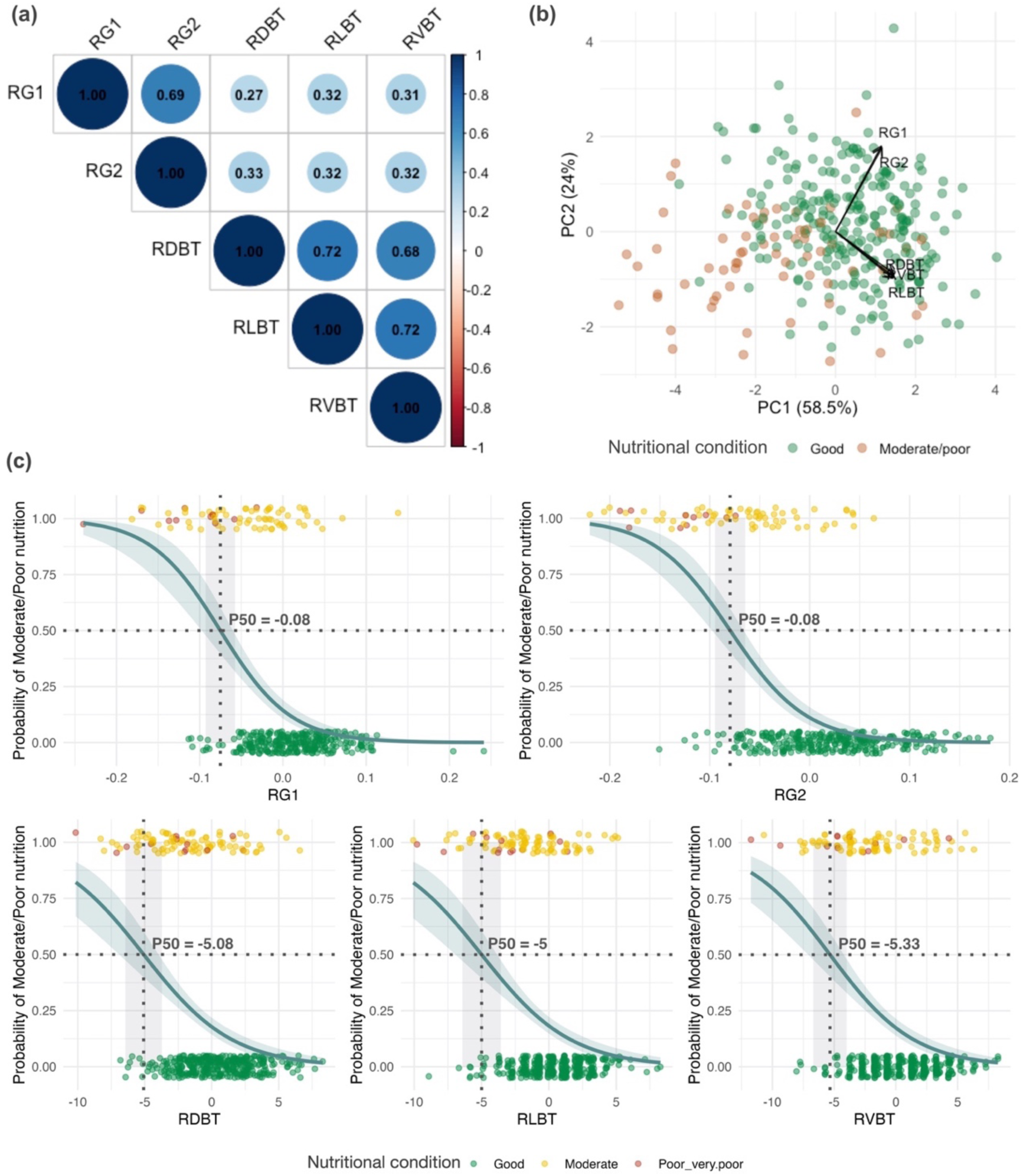
(a) Correlation matrix between all five Body Condition Indices (BCIs) (RG1, RG2, RDBT, RLBT, RVBT), showing Pearson correlation coefficients (α = 0.05). (b) Principal Component Analysis of the five BCIs. Points represent individual animals colored by nutritional condition (Good=green; Moderate/poor=orange). Arrows indicate the contribution of each BCI to the first two principal components, with axis labels showing the percentage of variance explained. (c) Logistic Generalised Linear Models (GLM) and their associated 95% CI, showing the probability of being in a moderate/poor nutritional condition according to each BCI value (RG1, RG2, RDBT, RLBT, RVBT). P50 represents the value of each BCI at which an individual has a 50% probability of being in a moderate/poor nutritional condition, effectively marking the midpoint of the logistic relationship between the BCIs and the risk of poor nutrition. Grey shading corresponds to the 95% CI of P50. Points represent raw data, jittered to facilitate visualization and the different colours indicate the nutritional condition of animals, which were clustered in two groups, i.e., animals in ‘good’ and ‘moderate/poor’ condition.

The PCA of the five BCIs showed that PC1 and PC2 explained 58.5% and 24% of the variance, respectively, capturing 82.5% of the total variation. We expected a reliable BCI to be high in animals with good nutritional condition and low in those with poor condition. Animals with good nutritional condition clustered in the top-right corner of the PCA, whereas those with moderate or poor condition grouped toward the bottom-left (Figure 3.b). PC1 captured overall variation in body condition, while PC2 distinguished two groups of BCIs. The two girth-based BCIs and the three blubber thickness-based BCIs each formed strongly correlated groups oriented in opposite directions along PC2, consistent with the Pearson correlation results. All five BCIs pointed toward the same side of the PCA, indicating that they generally reflected nutritional condition. However, RG1 and RG2 aligned with the top-right cluster of animals in good nutritional condition, whereas RDBT, RLBT, and RVBT pointed toward the bottom-right, overlapping with some animals of moderate or poor condition. Overall, the girth-based BCIs provided the clearest indicators of nutritional condition, while the blubber thickness-based BCIs were less discriminating, potentially influenced by other factors.

The five logistic GLMs fitted to assess the relationship between NCC and each BCI all showed overlap in BCI values between animals assigned to different NCCs (Figure 3.c), highlighting the subjectivity inherent in the assignment of the NCC. For each BCI, we estimated the value at which an individual had a 50% probability of being classified as moderate/poor (P50), representing the midpoint of the logistic relationship between the index and the probability of poor nutritional condition (see Table 1). Comparison of AIC values across the five models indicated that both girth-based indices were more strongly associated with nutritional condition than the three blubber thickness-based indices. Among the girth-based indices, RG2 was the most strongly associated with nutritional condition, while RVBT was the blubber thickness-based index most strongly related to the NCC (Table 1).

**Table 1.** Results of the logistic Generalised Linear Models (GLMs) showing parameters and coefficients for five separate models fitted to evaluate the relationship between nutritional condition (NCC) and each of the different body condition indices (BCIs) RG1, RG2, RDBT, RLBT, and RVBT. For each type of BCI (i.e., girth-based BCIs and blubber thickness-based BCI), the models with the lowest AIC are highlighted in bold. The total number of individuals included in each analysis is indicated in (N).

| Type of BCI | Model | Parameter | Estimate | SE | z value | AIC | P50 | N |
| --- | --- | --- | --- | --- | --- | --- | --- | --- |
| Girth-based BCIs | glm(NCC ~ RG1) | Intercept | -1.751 | 0.182 | -9.64 | 257.23 | -0.08 | 371 |
|  |  | RG1 | -23.546 | 3.484 | -6.76 |  |  |  |
|  | **glm(NCC ~ RG2)** | Intercept | -2.029 | 0.217 | -9.34 | **224.95** | -0.08 | 374 |
|  |  | RG2 | -25.218 | 3.361 | -7.50 |  |  |  |
| Blubber thickness-based BCIs | glm(NCC ~ RDBT) | Intercept | -1.564 | 0.161 | -9.72 | 296.77 | -5.08 | 481 |
|  |  | RDBT | -0.264 | 0.054 | -4.86 |  |  |  |
|  | glm(NCC ~ RLBT) | Intercept | -1.547 | 0.159 | -9.75 | 297.86 | -5 | 480 |
|  |  | RLBT | -0.280 | 0.058 | -4.79 |  |  |  |
|  | **glm(NCC ~ RVBT)** | Intercept | -1.666 | 0.172 | -9.66 | **281.13** | -5.33 | 480 |
|  |  | RVBT | -0.308 | 0.052 | -5.92 |  |  |  |

### 3.2. Temporal trends and confounding factors

The optimal model for RG2 retained the year with the sex interaction, COD, latitude, and decomposition stage as explanatory variables (Table S3). The main effect of the variable sex was not significant (p = 0.27), but sex was retained in the model due to the significant interaction with it in the year smoother. In males, there was a significant (p < 0.001) non-linear temporal trend in body condition, with minimum BCI values at the beginning of the time series in 1998, increasing to a peak in 2010, and subsequently declining through 2019. In females, no significant temporal trend was detected (p = 0.22) (Figure 4.a). COD was also retained in the model, but its effect was only marginally significant (p = 0.079), with slightly higher BCI values observed in individuals showing evidence of bycatch compared to those without. The effect of decomposition stages was also marginal (p= 0.093), with animals at a higher stage of decomposition (DCC3-4) showing slightly higher values than the fresher ones (DCC1-2). Latitude was retained in the model but its effect was not statistically significant (p = 0.21), with predicted RG2 values slightly declining with increasing latitude.

**Figure 4.**
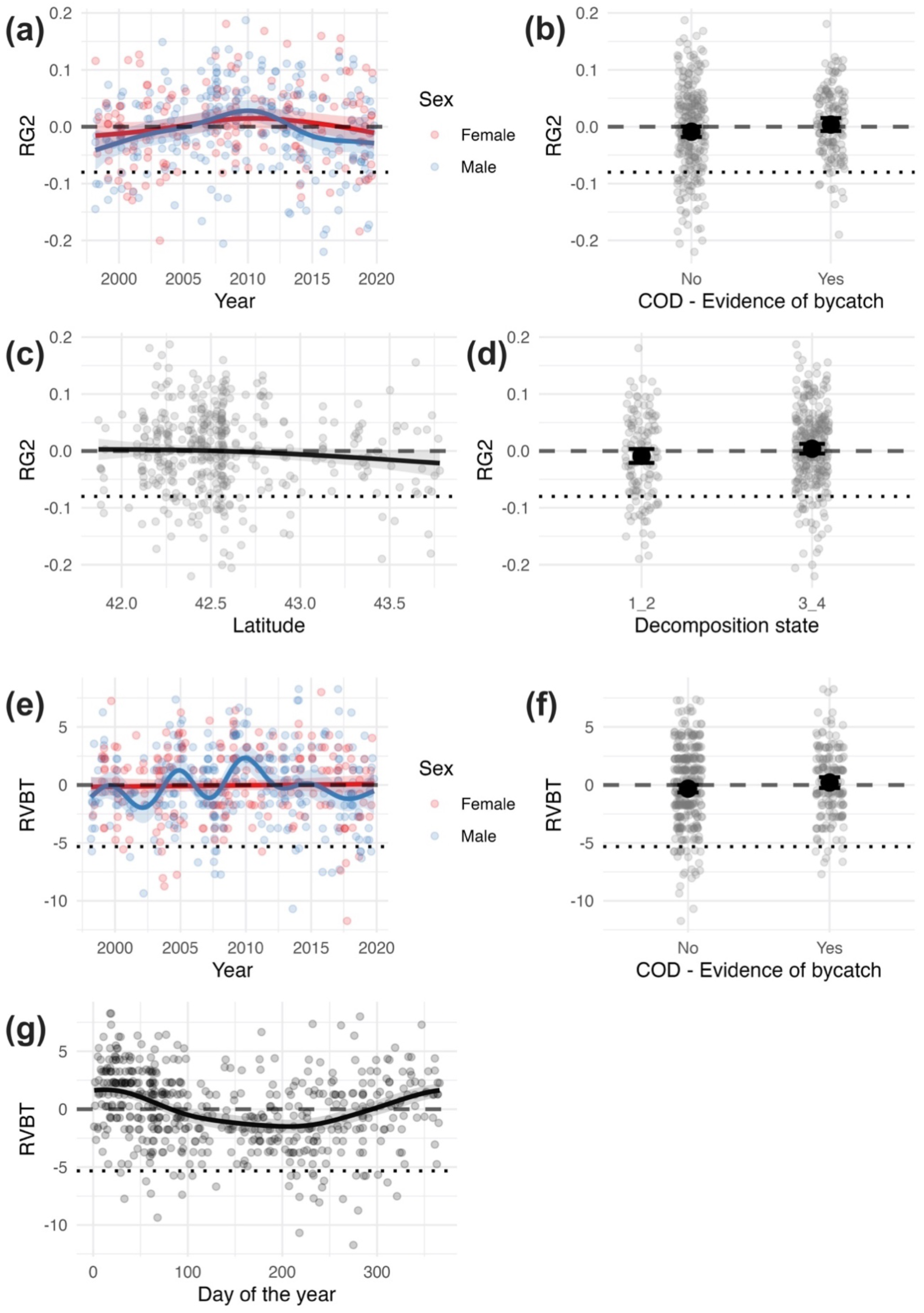
(a-d) Predicted RG2 values from the optimal GAM. Smooth terms (solid lines) and 95% confidence intervals (shaded area), showing (a) intraspecific temporal variation (1998-2019) separated by sex (males in blue, females in red), and (c) spatial variation per latitude. Overall differences between causes of death in relation to evidence of bycatch (b) and among decomposition stages (d), with black central points representing marginalized model estimates, and vertical error bars indicating the corresponding 95% confidence intervals. (e-g) Predicted RVBT values from the optimal GAM. Smooth terms (solid lines) and 95% confidence intervals (shaded area), showing intraspecific temporal variation over (e) the whole time period (1998-2019) separated by sex (males in blue, females in red), and (g) thorough the year. (f) Overall differences between causes of death in relation to evidence of bycatch, with black central points representing marginalized model estimates, and vertical error bars indicating the corresponding 95% confidence intervals. Semi-transparent raw data are overlaid in each plot. The dashed lines represent the average body condition (BCI=0) for the population, and the dotted lines represent the P50 value for RG2 (−0.08) and RVBT (−5.33) based on the NCC. The effect of the variable sex was not illustrated as it was not significant (and sex was retained in optimal models due to its significant interaction with year).

The optimal model for RVBT retained the year with the sex interaction, season, and COD (Table S3). The main effect of sex was not significant (p = 0.86), but sex was retained in the model due to its significant interaction with year. In males, there was a significant non-linear yearly trend in body condition (p < 0.001), with BCI values increasing to a peak in 2010, and subsequently declining through 2019 to attain near-initial values. There was no significant yearly trend in females (p=0.74) (Figure 4e). While the yearly trend for males showed higher variability than for RG2, the overall trend was similar. Predicted values of RVBT significantly varied across the year (p < 0.001) (Figure 4f). RVBT reached the highest values mid-January when SST were the lowest in Galicia (Figure S2), and the lowest values at the end of July, when water SST were higher (Figure S2). The COD was retained in the optimal model, but its effect was only marginally significant (p = 0.07), with slightly higher BCI values observed in individuals showing evidence of bycatch compared to those without.

As seen above in the GAM results, values of both BCIs differed between the two COD categories, even though the differences were only marginally statistically significant. Further exploration showed that animals with evidence of bycatch were on average in better nutritional condition than those without (81.1% *vs*. 64.5% in “good” condition for RG2 and 85.9% *vs*. 66.7% in “good” condition for RVBT) (Figure 5), although evidently the full range of nutritional status categories was seen within each COD category.

**Figure 5.**
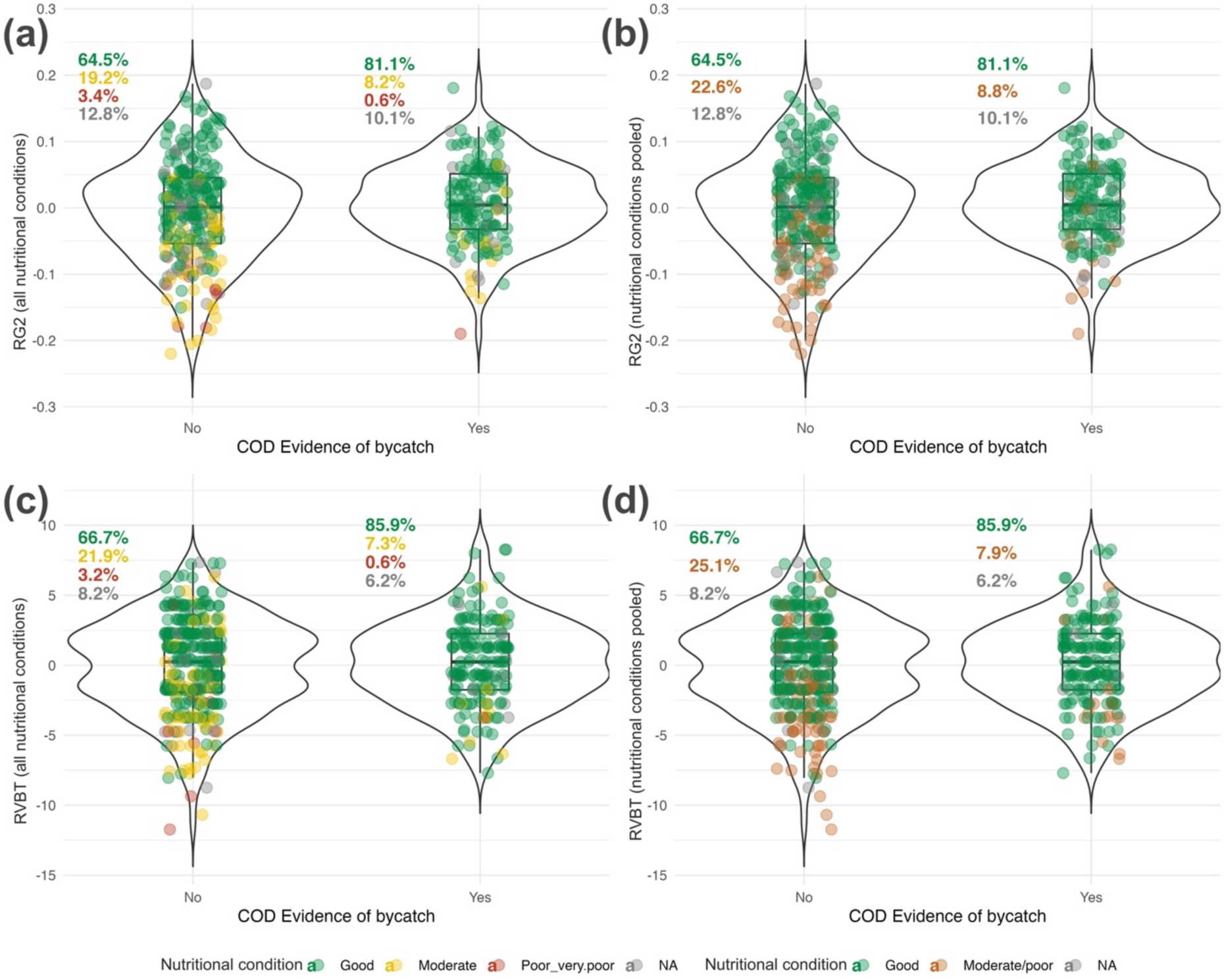
Violin plots showing the distribution of RVBT (top row) and RG2 (bottom row) in relation to the cause of death (COD) with regard to evidence of bycatch (‘No’ *vs*. ‘Yes’). (a) and (c) show all nutritional conditions categories (NCC) attributed during visual examination, while (b) and (d) show the categories ‘moderate’ and ‘poor/very poor’ pooled together. Individual points are colored according to NCC. In (a) and (c): Good (green), Moderate (yellow), Poor/very poor (red), NA (gray). In (b) and (d): Good (green), Moderate/poor (orange), NA (gray). The percentages displayed at the top-left of each violin indicate the proportion of animals within each NCC for that bycatch category, calculated relative to the total number of animals.

### 3.3. Trophic and environmental drivers

### 3.1. Temporal trends

The GAMs revealed a temporal trend in prey quality for male common dolphins between 1998 and 2018, with energy density values increasing significantly until 2006 and then declining thereafter. No temporal trend was detected in females (Figure 6a; Table S4). A significant temporal trend was identified for the DFA index based on prey SSB time series (p < 0.001; Table S4). Predicted values increased from 1998 to a peak in 2003, followed by a steady decline until 2013 and a subsequent recovery through 2018 (Figure 6h).

**Figure 6.**
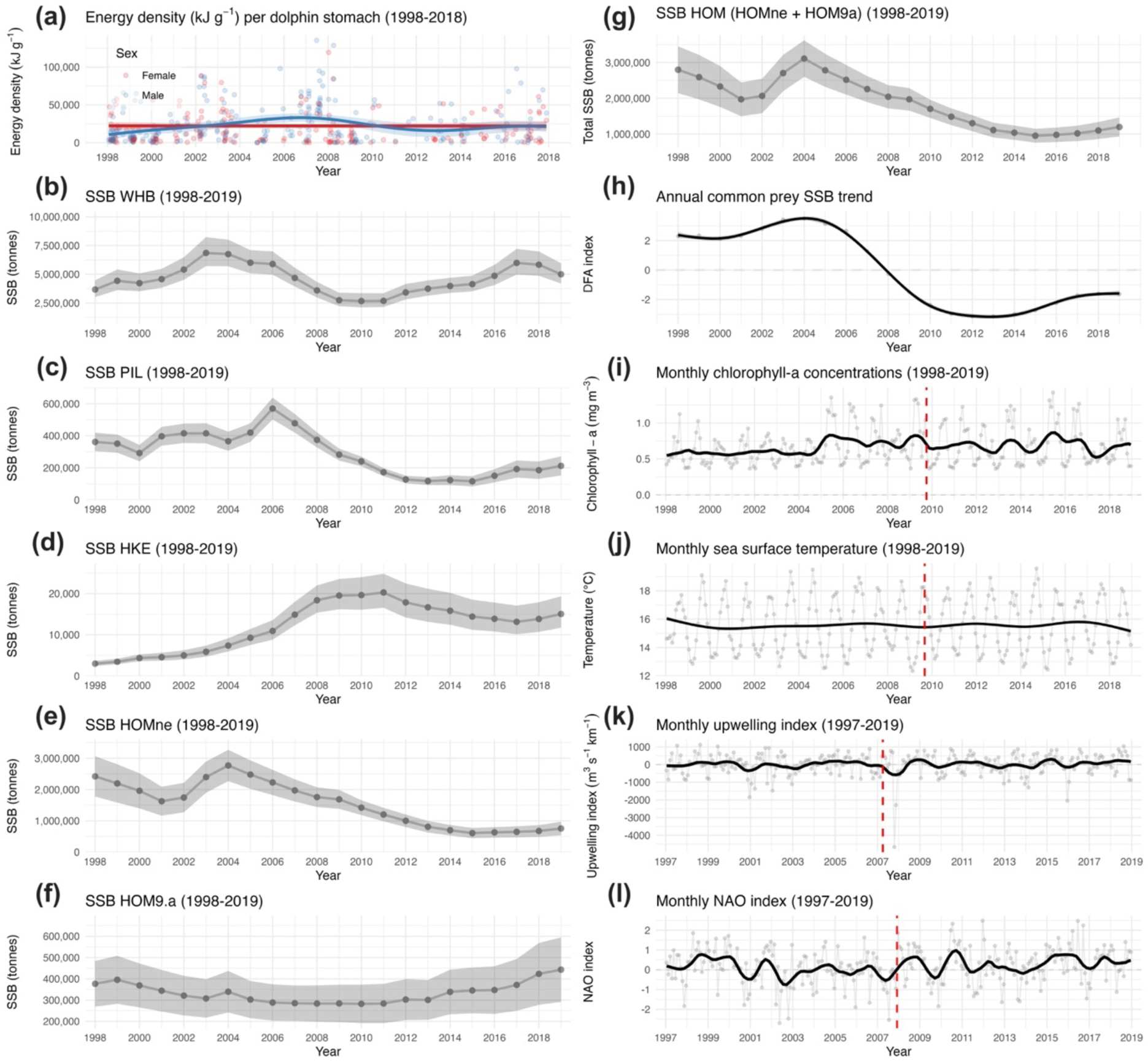
Temporal trends (1997-2019) in the trophic and environmental variables considered in this study in relation to common dolphin body condition. (a) Energy density per female (red) and male (blue) dolphin stomach, with GAM smooth terms (solid lines) and their 95% confidence intervals (shaded area), and semi-transparent raw data overlaid. (b) Annual Spanning Stock Biomass (SSB) values (grey points) of blue whiting in ICES subarea 27.1–9, 12 and 14, and their 95% confidence interval (shaded area). (c) Annual SSB values (grey points) of Iberian sardine in subarea 27. 8.c and 9.a, and their 95% confidence interval (shaded area). (d) Annual SSB values (grey points) of European hake in subarea 27. 8.c and 9.a, and their 95% confidence interval (shaded area). (e) Annual SSB values (grey points) of Atlantic horse mackerel in subarea 27.8 and divisions 2.a, 3.a, 4.a, 5.b, 6.a, 7.a–c, e– k, and their 95% confidence interval (shaded area). (f) Annual SSB values (grey points) of Atlantic horse mackerel in subarea 27.9.a, and their 95% confidence interval (shaded area). (g) Summed annual SSB values (grey points) of Atlantic horse mackerel in subarea 27.8 and divisions 2.a, 3.a, 4.a, 5.b, 6.a, 7.a–c, e–k, and subarea 27.9.a, and their 95% confidence interval (shaded area). (h) Annual DFA index based on prey SSB, summarizing the common trend in prey community composition, with GAM smooth terms (solid lines) and their 95% confidence intervals (shaded area). (i) Monthly mean of chlorophyll-a concentrations. (j) Monthly mean of sea surface temperature. (k) Monthly mean of upwelling intensity index. (l) Monthly mean of North Atlantic Oscillation index. In (i), (j), (k), and (l), the black line represents the de-seasonalized trend of the Seasonal-Trend Decomposition using Loess (STL) analysis, and the red dashed line indicates the primary breakpoint year.

Time series decomposition and structural change analysis revealed a significant shift in the NAO toward more positive phases across the study period (Mann-Kendall τ = 0.099, p = 0.014, Sen’s slope = 0.0094 units year^-1^) with a primary breakpoint detected in November 2007. This atmospheric forcing was reflected in the UI, which exhibited a highly significant long-term increase (τ = 0.15, p < 0.001) at a rate of approximately 4.83 m^3^ s^-1^ km^-1^ year^-1^, pivoting around a primary breakpoint in March 2007. While SST remained statistically stable over the long term (p = 0.429; Sen’s slope = 0.0021 °C year^-1^), a structural breakpoint was identified in August 2009. Parallel to these physical changes, CHL demonstrated a steady and significant rise (τ = 0.235, p < 0.001, Sen’s slope = 0.0047 mg m^-3^ year^-1^), with a primary breakpoint in September 2009. The proximity of the SST and CHL breakpoints (August–September 2009) following the 2007 atmospheric and physical shifts suggests a lagged ecosystem-wide regime shift. Additional secondary breakpoints were identified (see Table S5), though these represented lower-magnitude adjustments compared to the primary shifts of 2007-2009.

### 3.2. Structural Equation Modeling

For the RG2 subset, model comparison based on AIC showed that the inclusion of direct paths from environmental and seasonal drivers to dolphin body condition was statistically unsupported, with AIC increasing from 2194.2 for the mediated model to 2196.1 for SST_deco, 2195.5 for UI_deco, 2195.6 for NAO_deco, and 2195.5 for day of the year. On the other hand, the inclusion of direct paths from environmental and seasonal drivers to dolphin body condition in the RVBT model was supported: the AIC decreased from 4880.4 to 4824.2 when including the day of the year, 4864.8 for SST_deco, 4875.6 for UI_deco, and increased to 4882.2 for NAO_deco. Thus, the *a-priori* pSEM model was retained for RG2, while direct paths for day of the year and SST_deco in sub-model 4 were added in to the RVBT *a-priori* pSEM model (Table 2). A direct path for UI_deco was not added in the RVBT sub-model 4, due to its collinearity with SST_deco.

**Table 2.** Summary of the piecewise Structural Equation Model (pSEM) paths for the Galician marine ecosystem, for both body condition indices (BCIs). For each sub-model, model fit is summarized by edf / DF, the Crit. value (F-statistic for smooth terms, t-statistic for parametric terms, with higher values indicating a stronger signal-to-noise ratio), p-values for the specific paths (statistical significance indicated by an asterisk (*)), and R^2^ (proportion of variance in the Response variable explained by the combined Predictors at that specific path). NAO_deco (North Atlantic Oscillation, decomposed trend); UI_deco (Upwelling Index, decomposed trend); SST_deco (Sea Surface Temperature, decomposed trend); CHL_deco (Chlorophyll-a concentration, decomposed trend); Prey quantity (DFA index); prey quality (energy density per stomach); Day of the year (seasonality).

| BCI | Path (sub-model) | Response | Predictor | edf / DF | Crit. Value | P-value | R<sup>2</sup> |
| --- | --- | --- | --- | --- | --- | --- | --- |
| **RG2 (N=195)** | **(1) Atmospheric forcing** | UI_deco | NAO_deco | 7.22 | 23.60 | < 0.001* | 0.47 |
|  | **(2) Primary production** | CHL_deco | SST deco | 8.70 | 3.92 | < 0.001* | 0.51 |
|  |  |  | UI deco | 8.21 | 10.30 | < 0.001* |  |
|  | **(3) Intermediate trophic level** | Prey quantity (DFA) | CHL deco | 7.89 | 9.63 | < 0.001* | 0.59 |
|  |  |  | UI deco | 7.94 | 5.16 | < 0.001* |  |
|  |  |  | SST deco | 8.91 | 10.84 | < 0.001* |  |
|  | **(4) Higher predator** | Dolphin body condition (RG2) | Sex (Male) | 195.00 | -0.29 | 0.770 | 0.09 |
|  |  |  | DFA (Female) | 2.15 | 0.92 | 0.407 |  |
|  |  |  | DFA: (Male) | 5.50 | 2.15 | 0.046* |  |
|  |  |  | Energy density | 2.26 | 2.66 | 0.075 |  |
| **RVBT (N=254)** | **(1) Atmospheric forcing** | UI_deco | NAO_deco | 7.42 | 19.09 | < 0.001* | 0.35 |
|  | **(2) Primary production** | CHL_deco | SST deco | 8.45 | 3.74 | < 0.001* | 0.51 |
|  |  |  | UI deco | 8.25 | 15.75 | < 0.001* |  |
|  | **(3) Intermediate trophic level** | Prey quantity (DFA) | CHL deco | 8.06 | 13.95 | < 0.001* | 0.58 |
|  |  |  | UI deco | 8.93 | 6.57 | < 0.001* |  |
|  |  |  | SST deco | 8.95 | 11.98 | < 0.001* |  |
|  | **(4) Higher predator** | Dolphin body condition (RVBT) | DFA | 2.59 | 1.66 | 0.17 | 0.27 |
|  |  |  | Energy density | 1.00 | 0.15 | 0.69 |  |
|  |  |  | SST deco | 1.37 | 4.30 | 0.02* |  |
|  |  |  | Day of the year | 8.00 | 9.07 | < 0.001* |  |

Both RG2 and RVBT pSEM models achieved high explanatory power for environmental transitions (sub-model 1, 2, 3) (R^2^ = 0.35–0.59), with the intermediate prey quantity index (sub-model 3) respectively capturing 59% and 58% of the variance driven by combined oceanographic signals (Table 2). For RG2, the lower R^2^ observed in dolphin response suggests that ∼9% of the variance in body condition is explained by food-related variables. This suggests that while food consumption is a significant driver of body condition (p = 0.046 for males), other factors, such as diseases, are likely to be involved. The RG2 pSEM model results show that the Galician ecosystem operates as a strictly bottom-up mediated cascade, where climatic and oceanographic forcing does not directly impact dolphin body condition but must first propagate through the intermediate prey community. The RVBT pSEM model suggests that this BCI is strongly influenced by long-term variation in SST, and a seasonal component also likely related to SST (Table 2). None of the prey-related variables had a significant effect on this BCI (Figure 7). This result suggests that the RVBT BCI might not be suitable to investigate changes in prey composition and quality in environments with high seasonality in oceanographic parameters. Energy density had no significant effect on any BCIs within sub-model 4. However, the relationship for RG2 was near-significant (p = 0.075) and followed an expected biological trend, with body condition increasing with energy density before reaching a plateau at a maximum threshold (Figure 7).

**Figure 7.**
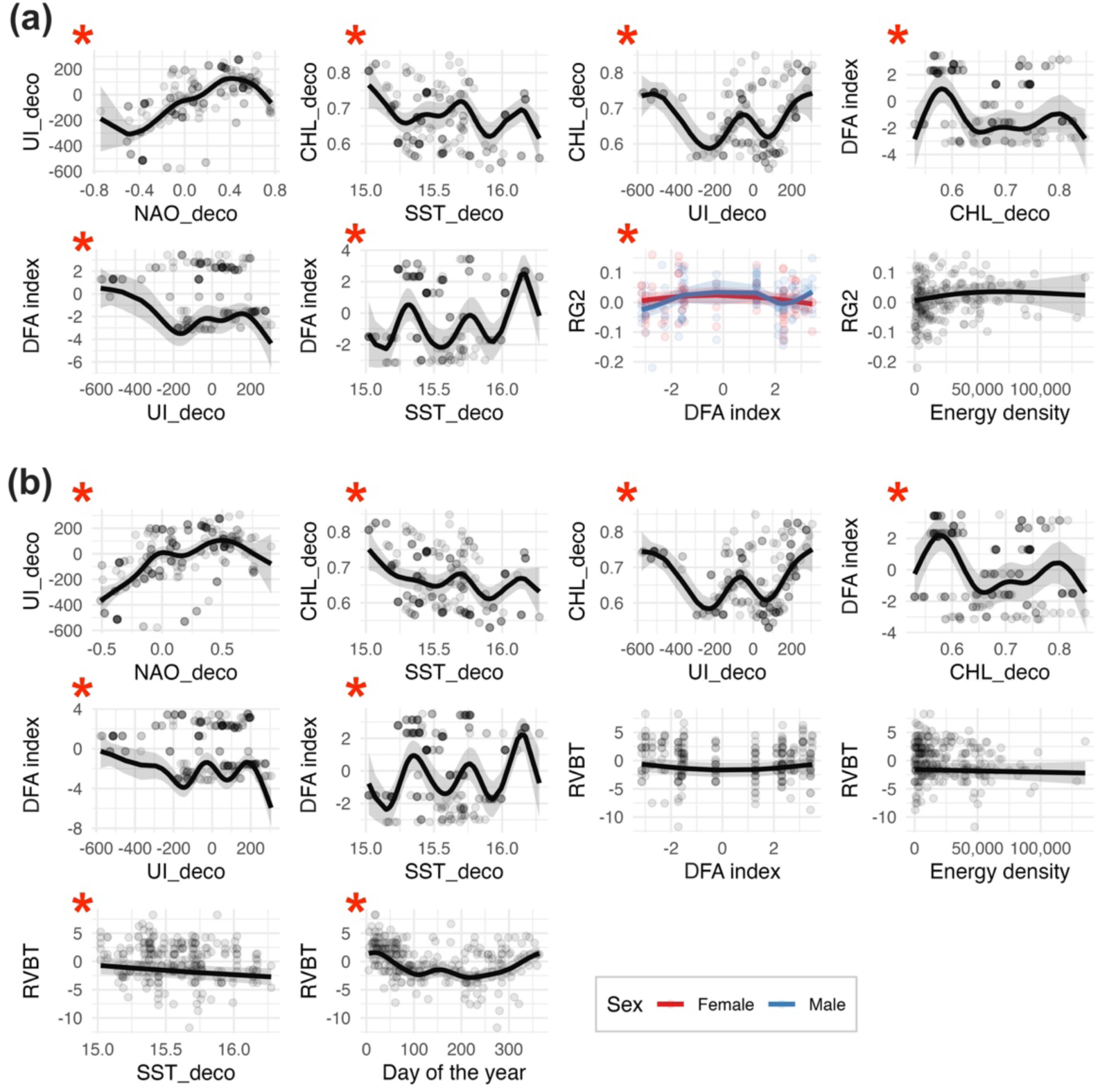
Visualization of the relationships between variables from the piecewise Structural Equation Model (pSEM) paths for the Galician marine ecosystem, for both body condition indices RG2 (a) and RVBT (b). Statistical significance between variables is indicated by a red asterisk (*). NAO_deco (North Atlantic Oscillation, decomposed trend); UI_deco (Upwelling Index, decomposed trend); SST_deco (Sea Surface Temperature, decomposed trend); CHL_deco (Chlorophyll-a concentration, decomposed trend); Prey quantity (DFA index).

For all sub-models, concurvity between variables remained within acceptable limits (< 0.65). Diagnostics tests for the environmental sub-models 1, 2, and 3 exhibited minor residual patterning (*DHARMa* KS-test: p < 0.05) attributed to the inherent heteroscedasticity and stochasticity of high-resolution oceanographic time-series. However, given the high F-statistics (e.g., F = 23.6 for the NAO_deco-UI_deco link in the RG2 pSEM sub-model 1), these sub-models were retained. The residual patterning represents high-frequency noise in environmental intensity rather than a failure to capture the primary, long-term ecological trends driving the system.

## 4. Discussion

### 4.1. The residuals approach as a reliable BCI

The choice of an appropriate BCI will depend on the assumptions underlying its calculation and the intended research objectives. Our residuals-based BCIs were specifically designed to minimise the effect of body length. We considered it advantageous to remove the length effect prior to analysis, to remove confounding effects of size, and because it facilitates the interpretation of the contribution of other explanatory variables.

The residuals-based BCIs are straightforward to compute from long-term stranding network datasets because they rely on routine morphometric measurements commonly collected during necropsies and field examinations (Petitguyot et al., 2025). We applied this methodology to five different measurements, although the same approach could be extended to derive BCIs from other morphometric variables. Because the residuals are calculated from a population-level model, they provide a measure of body condition relative to the population average and can be used to define thresholds for classifying body condition. They performed well in models to assess spatio-temporal trends in body condition, as well as the effects of season, decomposition stage, and cause of death. They also appear sensitive to trophic and environmental drivers, making them a useful tool for investigating ecosystem changes.

Because our results indicate a marginal effect of the COD on our indices, this suggests that our indices showed potential for differentiating animals with different “health status”. *A priori,* we would expect the health status of animals which died due to physical trauma to be more representative of the living population - hence on average they are expected to be healthier, when compared to animals which died due to other causes (typically including infectious diseases and other illnesses). In our dataset, non-bycaught individuals were more frequently in poor nutritional condition than bycaught animals (Figure 5), supporting the hypothesis that they were generally in poorer health. However, nutritional condition alone is not a reliable proxy for overall health, as animals in apparently good body condition may still suffer from severe pathological conditions (IJsseldijk et al., 2024; Lennon et al., 2025). Further validation using long-term pathological datasets is therefore needed, although the present results remain encouraging.

Among the tested indices, RG2 outperformed RVBT in predicting the nutritional condition code assigned at necropsy. However, this condition code is itself subjective and probably relies more on assessment of overall body shape than direct assessment of tissue reserves, which likely favours girth-based indices. The type of reference variable used for validation (e.g., NCC, BMI, or other indices) will inevitably influence which BCI appears most suitable, as these metrics do not necessarily capture identical aspects of condition. Despite this potential bias, our results are in accordance with Stepien et al. (2023), who found that the girth taken behind the pectoral fins (G2) was an excellent predictor of the BMI in harbour porpoises. In addition, body width measurements, which are analogous to girths, are commonly used to derive volume-based BCIs from unoccupied aerial vehicle photogrammetry (e.g., Christiansen et al., 2016) and have been shown to be good predictors of mass-based BCIs in bottlenose dolphins (*Tursiops* spp.) (Cicciarella et al., 2026). Furthermore, our girth- and blubber-based BCIs were only moderately correlated with each other and responded differently to biological and environmental drivers, suggesting that they capture complementary aspects of body condition. Although both were able to detect long-term changes, blubber thickness may be more strongly influenced by seasonality and environmental factors such as SST, whereas girth may better reflect changes in food intake and declines in muscle mass associated with prolonged starvation and muscle catabolism. This supports the use of RG2 or the combined use of both indices, particularly as blubber thickness remains a debated proxy of body condition in cetaceans (Kershaw et al., 2017; Derous et al., 2020).

Information from BCIs is highly relevant for European monitoring frameworks including assessments conducted under the Marine Strategy Framework Directive (MSFD; Directive 2008/56/EC) and the Convention for the Protection of the Marine Environment of the North-East Atlantic (OSPAR). The residuals-based BCIs approach could be applied for large-scale assessments at the ecoregion or management-unit level, by merging datasets from different stranding networks using identical morphometric measurements prior to deriving the BCIs. However, baseline body condition may vary among regions due to environmental and population-level differences, inherently complicating direct comparisons across areas regardless of what kind of BCI is used.

Further validation should compare residuals-based BCI with weight-based indices and alternative methods such as biomarkers-based approaches (see Castrillon and Bengtson Nash, 2020). Ultimately, the most reliable BCI is likely to be the one best suited to the available dataset and the specific research question.

### 4.2. Confounding factors affecting body condition

We assessed the potential confounding effect of decomposition on BCI values, as carcass decomposition can cause progressive bloating and blubber leakage, potentially altering soft tissues and biasing girth and blubber thickness measurements (Moore et al., 2020). DCC showed only a marginally significant effect, detected exclusively for the RG2 BCI, with higher DCCs associated with slightly higher RG2 values. Although this suggests that decomposition may introduce a minor bias, incorporating DCC into the model did not alter the significant temporal trend observed in male body condition. Overall, these findings suggest that it might be advantageous to select only relatively fresh carcasses to assess body condition but, as it might not always be possible, it highlights the importance of including DCC as a covariate in the models.

The strong seasonal pattern observed in RVBT revealed a decline in BCI values during the first part of the year, reaching the lowest levels during summer, and a subsequent increase until the end of the year. Similar seasonal trends have been reported in other small cetaceans, including common dolphins (Albrecht et al., 2024) and harbour porpoises (Kastelein et al., 2019; Siebert et al., 2022; Stepien et al., 2023). The summer decline coincides with warmer SSTs, which likely reduce the need for thermal insulation (Lockyer et al., 2003). It also overlaps with the reproductive period (Murphy et al., 2009), suggesting that individuals may mobilize fat reserves to meet the high energetic demands associated with reproduction. To compensate for this loss in blubber thickness, animals may increase energy intake from early autumn through late winter, potentially making them more vulnerable to the effects of fasting during this period (Kastelein et al., 2019; Gallagher et al., 2021). Seasonal changes in prey quality may also contribute to these patterns. Common dolphins in Galicia and elsewhere in the Northeast Atlantic exhibit marked seasonal variation in diet composition (Santos et al., 2013; Hernandez-Gonzalez et al., submitted), which could partly explain the observed variability in body condition, although further research is needed to confirm this relationship.

### 4.3. Drivers of spatio-temporal trends in body condition

Our results revealed no statistically significant spatial trend in BCI values, although latitude was retained in the RG2 model. The Galician coastline may be too limited to resolve spatial variation in a highly mobile pelagic species. Common dolphins can move over large spatial scales throughout the year in European Atlantic waters (Virgili et al., 2024), so it cannot be excluded that individuals that died in a given area may reflect nutritional conditions experienced elsewhere. In addition, stranding locations do not necessarily reflect mortality locations because carcasses can drift long distances under the influence of winds and currents (Peltier et al., 2012). Dolphins that strand in Galicia may have died in Portuguese waters, or further offshore in the northwestern waters of Spain (Saavedra et al., 2017; Deslias et al., 2024). Nonetheless, fresh or moderately decomposed animals are unlikely to have drifted for more than a few days, suggesting that death could have occurred near the stranding location.

We found a declining trend since 2010 in the body condition of male common dolphins in Galicia, which reached values below the threshold of the population average (BCI=0). Body condition in males was higher than female body condition between 2005 and 2013. Interpreting the consequences of this decline is challenging, as no threshold has yet been established in common dolphins for when reduced body condition begins to compromise health or survival. The condition threshold (P50) calculated suggested that this boundary was crossed intermittently throughout the analyzed time period, but its reliability is limited since this threshold relies on the NCC. Further work is needed to fully understand the consequence of crossing these two types of thresholds we used in this study. Despite this, the decline in the body condition of males is potentially concerning and aligns with broader trends across the Northeast Atlantic. Similar declines in body condition have been reported in common dolphins in the Celtic Seas, where poorer condition was linked to increased starvation-related strandings in the UK and Ireland between 1990 and 2019 (Albrecht et al., 2024). Comparable patterns have also been observed in other marine mammals elsewhere, such as in the Baltic Sea, with declines in body condition of grey seals (*Halichoerus grypus*) between 2002 and 2015 (Kauhala et al., 2017), harbour seals (*Phoca vitulina*) between 2002 and 2016 and harbour porpoises from 1990 to 2016 (Siebert et al., 2022). Together, these findings point to a widespread deterioration in marine predator health.

Based on the best BCI to predict nutritional status (RG2), our results suggest that the declining body condition of male common dolphins in Galicia is likely driven by a bottom-up cascade, where climate and oceanographic variations alter prey community composition and reduce the quality of prey available to dolphins. A critical transition period was identified between 2007 and 2009, marked by an intensification of the NAO and UI, and an increase in CHL. Although no significant long-term trend in SST was detected, we identified a breakpoint around the same period, suggesting that more subtle shifts in SST may still have occurred. Such regime shifts likely disrupted the availability and distribution of the primary prey species of common dolphins. Fluctuations in the abundance of the four primary prey stocks for common dolphins in Galicia have been linked to fishing pressure alongside biological and environmental drivers, including the NAO, UI, SST, and CHL (Carrera and Porteiro, 2003; Cabrero et al., 2019; Pennino et al., 2019; Zimmermann and Werner, 2019; Izquierdo et al., 2021; Ferreira et al., 2023). However, the precise mechanistic pathways linking these environmental parameters to specific prey abundance remain to be fully elucidated.

Throughout the study period, there was a profound structural shift in the fish community, at least in species eaten by the dolphins. Between 1998 and 2005, the system was dominated by high-calorie, lipid-rich schooling prey (sardines, horse mackerel, blue whiting), likely explaining the elevated dolphin body condition observed through 2009. Conversely, from 2005 to 2013, these species progressively declined as the system transitioned to lower-calorie demersal prey, such as hake. This low-caloric regime persisted until approximately 2019, directly mirroring the post-2009 decline in male dolphin body condition. Temporal changes in prey composition mirrored shifts in the energy density consumed by male dolphins, as suggested by our dietary data. However, peaks in energy intake preceded peaks in body condition by 2-3 years (Figure 7). This lag may reflect limited overlap between datasets (Figure S3), as well as methodological constraints, including the use of fixed species-specific energy density values that do not account for potential spatio-temporal variation in prey lipid content. In addition, stomach contents reflect short-term dietary intake, whereas body condition may integrate physiological responses over longer time scales. It is unknown how rapidly common dolphins lose blubber thickness. Harbour porpoises can lose 0-3 mm of blubber thickness and 0.7-3 kg of body mass daily under near-fasting conditions (Kastelein et al., 2019). While rapid at the individual level, such changes may not immediately translate into population-level trends.

The temporal decline in energy density per male dolphin stomach observed in our data is not an isolated regional issue. Between 2002 and 2022, significant declines in fish body condition were documented in the Bay of Biscay and the Celtic Seas (Gernez et al., 2025), including prey targeted by common dolphins in Galicia, the Bay of Biscay and Celtic Seas, i.e., sardines, anchovies and hakes (Santos et al., 2013; Faure et al., 2025; Albrecht et al., 2026; Hernandez-Gonzalez, submitted). In the Bay of Biscay, a severe decline in the energy content of prey species of common dolphins have been documented, primarily driven by a decrease in the length and energy density of these species. Alarmingly, it is estimated that common dolphins in that region now have access to half the energy per individual prey item compared to 20 years ago (Favreau et al., 2025). This nutritional deficit is cascading upward from lower trophic levels; there appears to be a parallel decline in the quality of some of the prey consumed by the dolphins’ prey. For instance, comparisons of fish sampled in the Bay of Biscay during 2002-2008 and 2022-2023 showed significant declines in nutritional quality across multiple species, including small pelagic fish that serve as key prey for hake and whiting (Amelot et al., 2025). The authors attributed these declines to changes in the quantity and/or quality of phytoplankton and zooplankton resources at the base of the food web. A similar mechanism may be operating in Galicia. Climate- and hydrography-driven regime shifts in phyto- and zooplankton communities occurred around 1997-1998, marking a transition from cold, dry to warm, wet conditions (Bode et al., 2020). Such changes could have altered the dynamics of the transfer of energy through the foodweb.

The lack of temporal changes in body condition and energy density of prey consumed by female common dolphins could be linked to sexual segregation within this population (Meynier et al., 2008; Viricel et al., 2008; Fernández-Contreras et al., 2010; Ball et al., 2017), whereby males and females exhibit distinct habitat-use patterns and dietary preferences. Such differences could influence exposure to temporal variation in prey resources. Nevertheless, additional research is required to better understand the mechanisms driving these observed patterns.

While environmental and trophic variables explained a substantial portion of the variation in dolphin body condition in our dataset, the relatively low deviance explained by our pSEM models suggests that additional unmeasured factors contribute to the long-term decline in the body condition indices. Some of this unexplained variation likely reflects confounding factors discussed previously (DDC and COD). Health-related processes may also play an important role. Pathogens, parasite burdens, and other pathological conditions could interact synergistically with nutritional stress, reducing body condition in ways not captured by trophic and environmental indices alone. For example, both the prevalence of *Anisakis* spp. in the stomachs of stranded common dolphins in Galicia and the occurrence of associated gastric ulcers have increased in recent decades (Pons-Bordas et al., 2020; Rueda-Díez et al., 2026). Integrating health indicators may therefore improve future model performance and predictive power.

## 5. Conclusions

Our results uncovered a decline in body condition of common dolphins in Galician waters over the past decades, which was likely driven, at least in part, by bottom-up ecological processes, whereby climate and oceanographic changes alter prey community composition and reduce prey quality available to dolphins. These findings add to growing evidence that both prey quantity and quality strongly influence the body condition and nutritional status of marine mammals, ultimately affecting their health, reproductive success, and survival (e.g., Kauhala et al., 2018; Castrillon and Bengtson Nash, 2020; IJsseldijk et al., 2021). Nevertheless, the observed decline is likely multifactorial and may also reflect the influence of diseases and other health-related stressors acting alongside environmental changes.

Our study demonstrates that residual-BCIs derived from morphometric measurements effectively capture variation in body condition in common dolphins, and this approach is likely applicable to other species. In particular, girth-based BCIs appeared more robust than blubber thickness-based indices in environments characterized by strong seasonal variability. We also highlighted the importance of accounting for potential confounding factors when assessing body condition in small cetaceans. These BCIs can contribute to an integrated framework for detecting subtle physiological changes and evaluating how ecological variability may affect population health over time.

Declining body condition in marine predators and their prey has been documented in several regions worldwide and is expected to intensify as climate change continues to disrupt marine ecosystems. In the Northeast Atlantic, ongoing environmental changes are likely to further affect the prey resources of common dolphins. Future projections suggest that the species may shift its distribution northward in response to these changes (Lambert et al., 2011; Chandelier and Kiszka, 2026), potentially with additional consequences for nutritional status, health, and survival.

In this context, the development and application of reliable BCIs are becoming increasingly important for tracking long-term ecosystem change. We believe the indices proposed here could provide a standardized approach for large-scale assessments under regional and European monitoring frameworks such as OSPAR and the MSFD. Such assessments rely heavily on the long-term collection of biological data and samples by stranding networks – not only sex, length, girth and blubber thickness but also age, reproductive status, and health (as revealed by pathological, histopathological and ecotoxicological analyses), emphasizing the critical need for sustained and adequate funding to support these programs (Petitguyot et al., 2025). More broadly, our findings underscore the value of integrated, long-term monitoring frameworks combining ecological, pathological, and physiological indicators to better identify emerging pressures and support adaptive management of cetacean populations in the region.

## CRediT authorship contribution statement

Marie A.C. Petitguyot: Writing – review & editing, Writing – original draft, Methodology, Formal statistical analysis, Data curation, Conceptualization.

Juliette Champsaur: Writing – review & editing, Formal statistical analysis, Data curation. Alberto Hernandez-Gonzalez: Writing – review & editing, Formal analysis, Data curation.

Alfredo López, Pablo Covelo, Jose Martínez-Cedeira, Xabier Pin, Mónica González, Uxía Vázquez: Writing – review & editing, Data curation, Resources, Funding acquisition.

Graham J. Pierce: Writing – review & editing, Methodology, Conceptualization, Supervision, Funding acquisition.

## Acknowledgments

We thank Antonio Bode at the IEO-CSIC Coruña (Spain) for providing us with upwelling index data. This research was funded under the SeaChanges ITN - Marie Skłodowska-Curie Actions-ITN grant n°813383, the CetAMBICion project and (Co-ordinated strategy for the assessment, monitoring and management of cetaceans in the Bay of Biscay and Iberian coast subregion; EU, DG-ENV/MSFD 2020), the EMPHATIC project (E-DNA, Microbiomes, Photogrammetry and Hormones - Assessment Techniques In Cetaceans, Biodiversa+ BioDivMon), the MERMA CIFRA project (Monitorización, Evaluación y Reducción de la Mortalidad Accidental de Cetáceos debido a Interacciones con la Flota Española: Revisión y Acción, Ministerio de Agricultura, Pesca y Alimentacion (Gobierno de España), MAP2021-02) and the CIBBRiNA project (Coordinated Development and Implementation of Best Practice in Bycatch Reduction in the North Atlantic, Baltic and Mediterranean Regions; LIFE22-NAT-NL-LIFE-CIBBRiNA/101114301). The stranding network in Galicia is managed by the Coordinadora para o Estudo dos Mamiferos Mariños (CEMMA) and has the support of the Xunta de Galicia/Dirección General de Patrimonio Natural.

## Data Availability Statement

The datasets and R code for this study are available here: https://doi.org/10.5281/zenodo.21378601. All environmental data was obtained from publicly accessible online repositories.

