## Supplementary material for "Body condition as an indicator of long-term ecosystem change: evidence from common dolphins"

**Table S1.** Definition of nutritional condition state (NCC) categories used in this study, adapted from Kuiken and García-Hartmann (1991).

| NCC | Definition |
|---|---|
| Good | Well-developed back musculature; a convex profile; a high proportion of body fat. |
| Moderate | Normal development of back musculature; a straight profile; an elongated body appearance, with little body fat. |
| Poor/very poor | More or less depressed back musculature; a concave profile; ribs, scapula, or vertebral processes may be visible; very reduced or practically nonexistent body fat; serous atrophy of the fat is observed. |

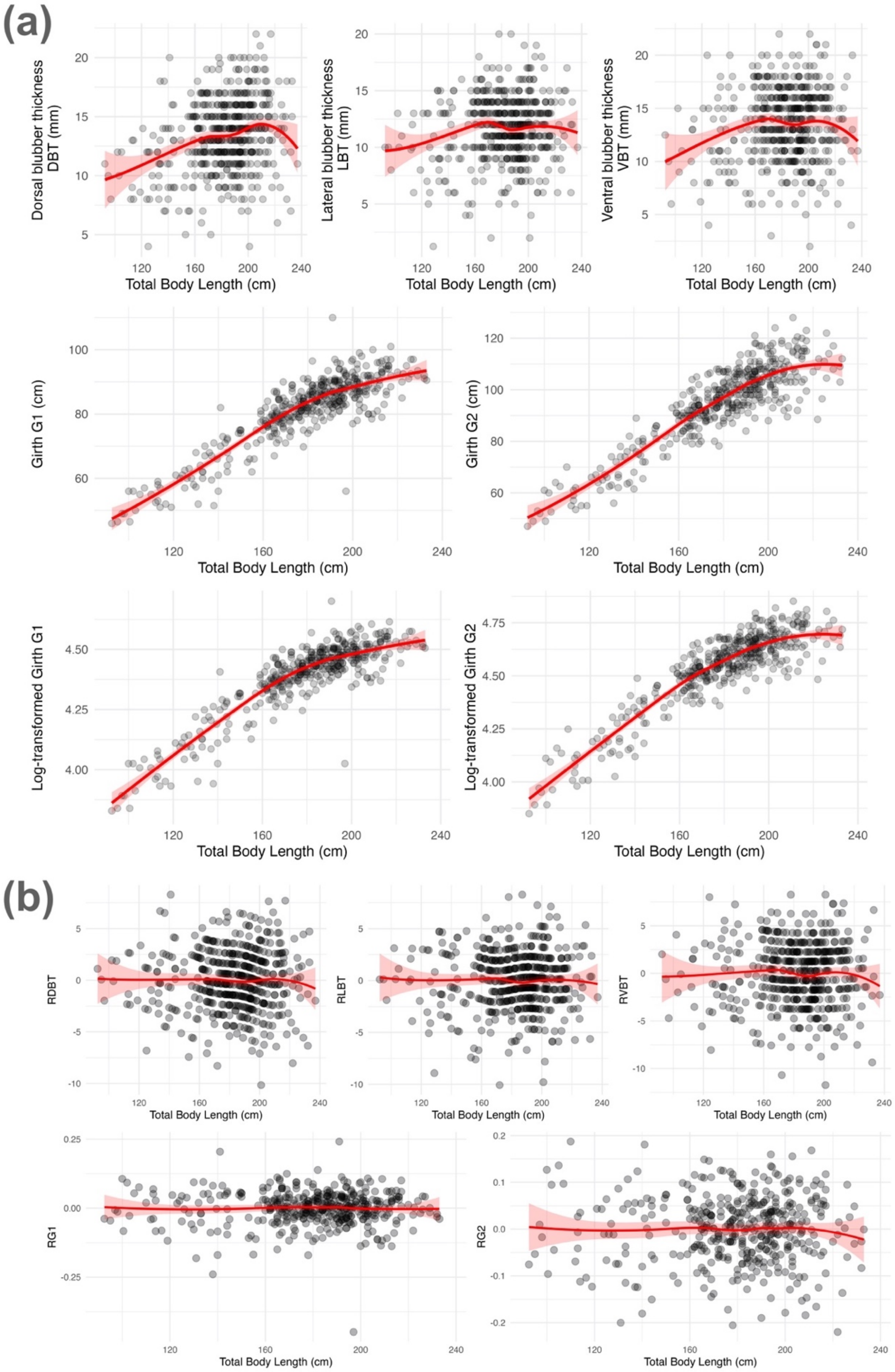


**Figure S1.** Relationship between (a) body condition metrics, i.e., the dorsal blubber thickness (DBT), lateral blubber thickness (LBT), ventral blubber thickness (VBT), the girth taken at the post-nuchal depression (G1), the girth taken behind the pectoral fins (G2), and total body length, showing non-linearity and heteroscedasticity; and between (b) the computed residual-BCIs and total body length. In each figure, the black dots represent individual data points, while the red curve depicts a fitted smooth regression line with its associated 95% confidence interval.

**Table S2.** Results of the Generalized Additive Models (GAMs) explaining variation in morphometric measurements taken on dead common dolphins (G1: girth taken at the post-nuchal depression, G2: girth taken after the pectoral fins, dorsal blubber thickness, lateral blubber thickness, ventral blubber thickness) in function of the total body length (L1) and the sex. Models with or without an interaction between the total body length and the sex are compared with the Akaike Information Criterion (AIC). The total number of individuals included in each analysis (N) is presented.

| **Morphometric measure** | **Explanatory variables in GAM** | **Results** | **AIC** | **N** |
|---|---|---|---|---|
| (1) log(G1) | ~ s(Body length) + Sex | s(L1): p=<2e-16 ***; Sex: p=0.39 | -1135.61 | 421 |
| | ~ s(Body length, by=Sex) + Sex | s(L1): for both males and females p=<2e-16 ***; Sex: p=0.93 | -1137.53 | |
| (2) log(G2) | ~ s(Body length) + Sex | s(L1): p=<2e-16 ***; Sex: p=0.38 | -1024.76 | 424 |
| | ~ s(Body length, by=Sex) + Sex | s(L1): for both males and females p=<2e-16 ***; Sex: p=0.42 | -1023.07 | |
| (3) Dorsal blubber thickness | ~ s(Body length) + Sex | s(L1): p=<2e-16 ***; Sex: p=0.45 | 2603.04 | 520 |
| | ~ s(Body length, by=Sex) + Sex | s(L1): Females p=0.0007 ***, males p=5.28e-07 ***; Sex: p= 0.35 | 2603.37 | |
| (4) Lateral blubber thickness | ~ s(Body length) + Sex | s(L1): p=0.00718 **; Sex: p=0.84 | 2540.29 | 520 |
| | ~ s(Body length, by=Sex) + Sex | s(L1): females p= 0.439, males p=0.0013 **; Sex: p=0.89 | 2539.02 | |
| (5) Ventral blubber thickness | ~ s(Body length) + Sex | s(L1): p=0.003 **; Sex: p=0.55 | 2707.98 | 519 |
| | ~ s(Body length, by=Sex) + Sex | s(L1): females p= 0.50, males p=0.002 **; Sex: p=0.47 | 2707.85 | |

**Table S3**. Results of the Generalized Additive Models (GAMs) explaining variation in values of body condition indices of common dolphins (RG2 and RVBT) in function of spatio-temporal variables and confounding factors (i.e., time (yeardec), season (day of the year), cause of death with respect to bycatch (COD_bycatch), decomposition stage (DCC), and space (latitude)). For each response variable, both the full model and the optimal model selected via backward stepwise Akaike Information Criterion (AIC) procedure are presented. Model goodness of fit is summarized by $R^2$, deviance explained (%DE) and AIC. The total number of individuals included in each analysis (N) is also shown.

| BCI | Model | $R^2$ | %DE | AIC | N |
|---|---|---|---|---|---|
| RG2 | Full model: ~ s(yeardec, by=Sex) + s(Day of the year, bs="cc") + Sex + COD_bycatch + DCC + s(Latitude) | 0.074 | 9.96 | -1046.63 | 421 |
| | Optimal model: ~ s(yeardec, by=Sex) + Sex + COD_bycatch + DCC + s(Latitude) | 0.073 | 9.78 | -1046.57 | |
| RVBT | Full model: ~ s(yeardec, by=Sex, k=11) + s(Day_of_year, bs="cc") + Sex + COD_bycatch + DCC + s(Latitude) | 0.146 | 16.6 | 2611.24 | 515 |
| | Optimal model: ~ s(yeardec, by=Sex, k=11) + s(Day_of_year, bs="cc") + Sex + COD_bycatch | 0.178 | 20.3 | 2595.93 | |

**Table S4.** Results of the Generalized Additive Models (GAMs) explaining temporal trends in proxies for prey quality and prey availability between 1998-2019. For each response variable, model fit is summarized by the p value, $R^2$, deviance explained (%DE), AIC, and the total number of individuals included in each analysis (N).

| Trophic / environmental driver (period) | Proxy | Model | P value | R2 | %DE | AIC | N |
|---|---|---|---|---|---|---|---|
| Prey quality (1998-2018) | Energy density per stomach | Energy_kJ ~ s(yeardec, by = Sex) | < 0.01 | 0.04 | 5.69 | 13477.58 | 448 |
| Prey availability (1998-2019) | DFA index based on SSBs of common dolphin main prey | Prey_DFA ~ s(Year) | < 0.001 | 0.998 | 99.9 | -29.09 | 22 |

**Table S5.** Results of the structural change analysis (1997-2019) and identification of breakpoints years. For each environmental variable, the time period, the Sen's slope, the Mann-Kendall test (MK τ), p-value, primary and secondary breakpoints years, and the overall trend, is identified.

| **Variable** | **Period** | **Annual Sen's slope** | **MK τ** | **p-value** | **Primary breakpoint** | **Secondary breakpoints** | **Trend result** |
|---|---|---|---|---|---|---|---|
| **NAO Index** | 1997-2019 | +0.0094 | 0.099 | **0.014** | **Nov 2007** | 2000, 2011, 2014 | Significant increase |
| **Upwelling Index** | 1997-2019 | +4.8257 | 0.150 | **< 0.001** | **March 2007** | 2000, 2003, 2014 | Significant increase |
| **SST** | 1998-2019 | +0.0021 | 0.032 | 0.429 | **Aug 2009** | 2001, 2006, 2012 | Stable (buffered) |
| **Chlorophyll-a** | 1998-2019 | +0.0048 | 0.235 | **< 0.001** | **Sept 2009** | 2004, 2013, 2016 | Significant increase |

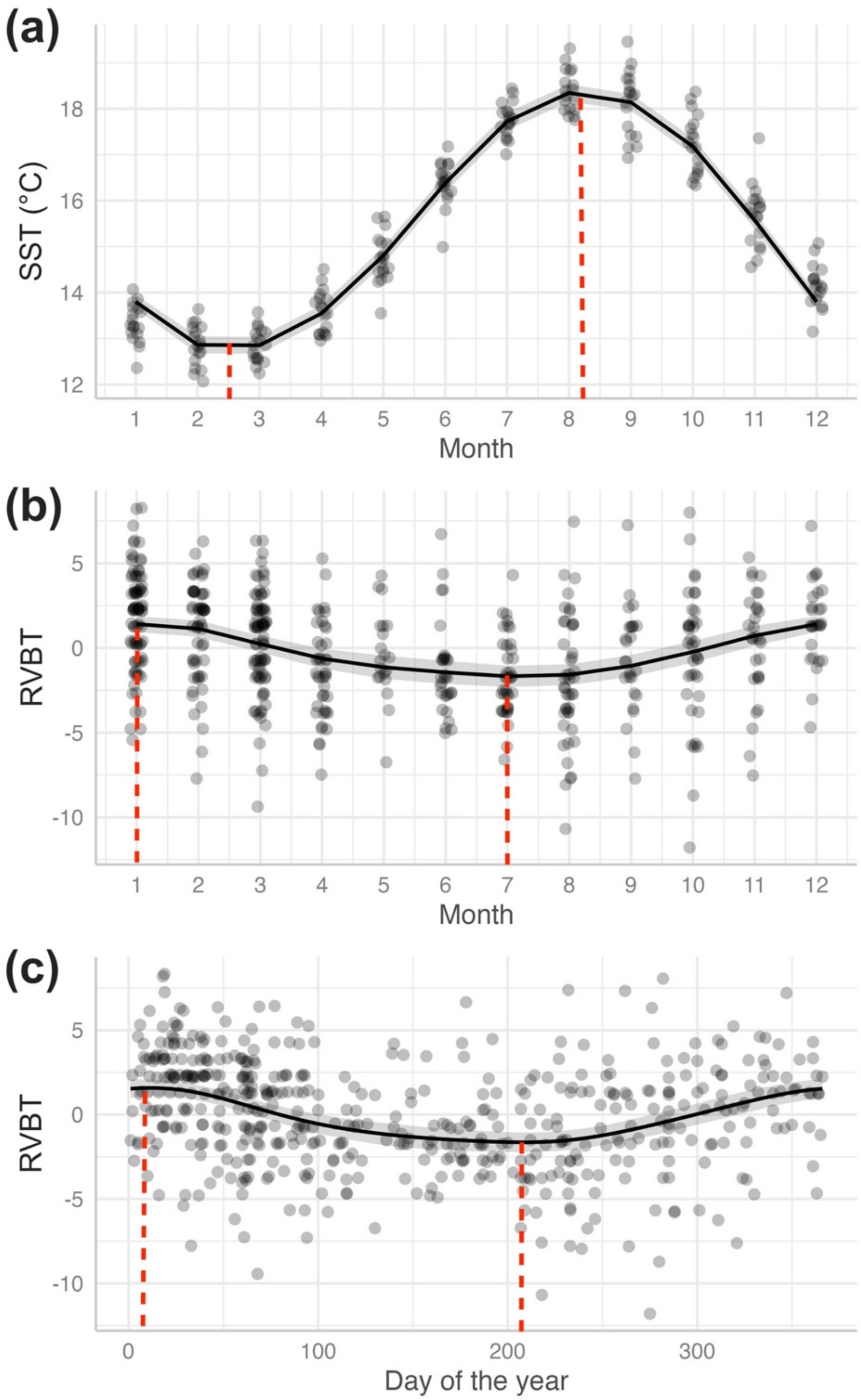


**Figure S2**. Seasonal trends in sea surface temperature (°C) (a) and ventral blubber based-BCI (RVBT) visualized by month (b) and by day of the year (c).

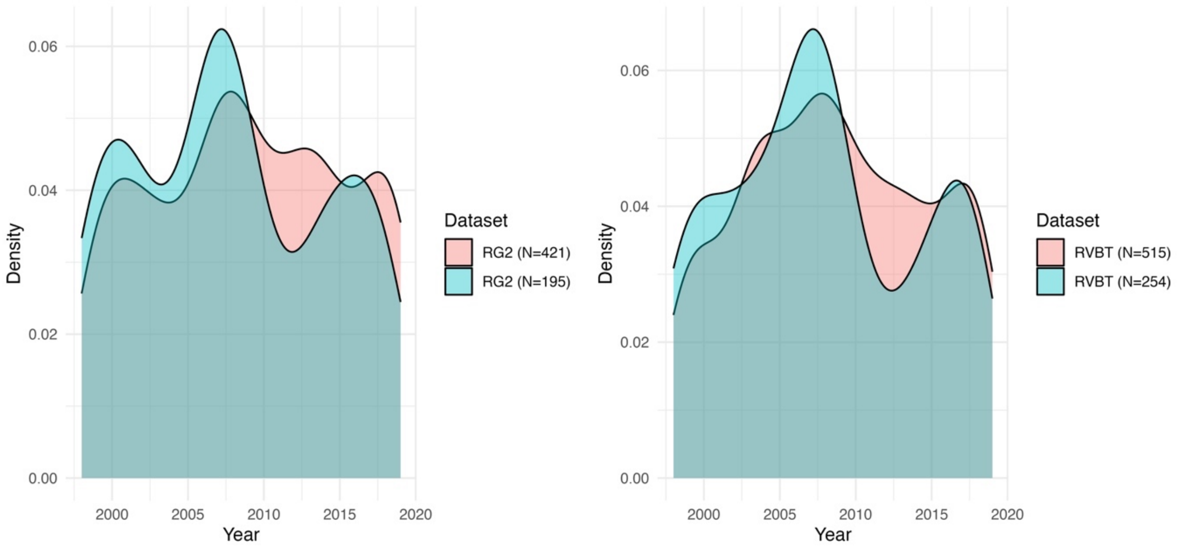


**Figure S3.** Density of dolphins per year in the datasets of RG2 and RVBT, showing the reduced number of animals present between the years 2010-2015 in the smaller subsets of both body condition indices (in blue) compared to the larger datasets (in red).
